# Super Pangenome of Grapevines Empowers Improvement of the Oldest Domesticated Fruit

**DOI:** 10.1101/2024.02.28.582440

**Authors:** Li Guo, Xiangfeng Wang, Dilay Hazal Ayhan, Mohammad Saidur Rhaman, Ming Yan, Jianfu Jiang, Dongyue Wang, Wei Zheng, Junjie Mei, Wei Ji, Jian Jiao, Shaoyin Chen, Jie Sun, Shu Yi, Dian Meng, Jing Wang, Mohammad Nasim Bhuiyan, Guochen Qin, Linling Guo, Qingxian Yang, Xuenan Zhang, Haisheng Sun, Chonghuai Liu, Wenxiu Ye

## Abstract

Grapevine (*Vitis*) is the oldest domesticated fruit crop with great cultural and economic importance. Here, we assemble and annotate haplotype-resolved genomes of 72 *Vitis* accessions including 25 wild and 47 cultivated grapevines, and a haplotype-resolved complete genome of *V. vinifera*. Coalescent phylogenomics of 142 haplotype genomes disentangles the mysterious hybridization history of grapevines, revealing enormous genetic diversity among species. Pangenome analysis together with phenotyping data reveals that European cultivars, more susceptible to the most destructive disease downy mildew (DM), had a smaller repertoire of disease resistance genes of NLR family. Through extensive structural variation (SV) characterization, phenotyping, transcriptome profiling of 113 *Vitis* accessions, and SV-eQTL analysis, we have identified over 79 SVs and their relevant genes significantly associated with DM resistance, exemplified by a lysine histidine transporter, *VvLHT8*. This haplotype-resolved complete genome and pangenome of *Vitis* genus will accelerate grapevine breeding and enrich our understanding of the evolution and biology of grapevines.

## Main

Grapevine (*Vitis*) is one of the oldest domesticated (∼11,000 BC) crops with great influence over human civilization^1^. Since grapes, the berries of grapevines, are economically important table fruits and ingredients for many commodities such as wine, juice, and vinegar, there is a constant interest in improving the grapevine germplasms in terms of berry quality, resistance to biotic and abiotic stresses, convenience for cultivation. However, years of continuous domestication and breeding make modern grapevine cultivars quite narrow in genetic diversity^2–4^ and vulnerable to a variety of stresses such as cold, downy mildew, and grey mold, posing a threat to viticulture and the entire grape industry. The wild grapevine accessions harbor critical genetic diversity vital to improving cultivated grapevines, but so far have not been comprehensively investigated and sufficiently utilized in breeding. The *Vitis* genus hosts two subgenera, *Muscadinia* and *Euvitis* with roughly 70 inter-fertile wild species mainly categorized into three clades, European, North American and East Asian grapevine, based on their geographical distribution^5,6^. The common European grapevine, *V. vinifera*, has a long history of domestication with thousands of commercial cultivars for fresh consumption and wine making, while several North American species such as *V. labrusaca*, *V. riparia*, *V. berlandieri*, have also been used in interspecific breeding program mainly for making stress-resistant cultivars and rootstocks. In contrast, the Asian species are much less explored for breeding but have the largest number of species thus genetic diversity^5^.

High-quality genomic information lays the foundation for modern crop breeding approaches and genetic studies. Although a highly homozygous grapevine *V. vinifera* cv. PN40024 was once the fourth plant to have its genome sequence assembled^7^, *Vitis* genomics fell much behind other crops with several highly fragmented genomes^8–11^. Until recently a gap-free PN40024 genome assembly was reported^12^, but it provided limited insights to the clonally propagated crops of permanent heterozygosity such as grape cultivars^9^. In fact, it still contains several heterozygous loci with two haplotypes collapsed in a diploid assembly suffering from assembly errors. Given that grapevine genomes are highly heterozygous due to frequent outbreeding, a haplotype-aware assembly is not only required for assembly accuracy, but also critical for understanding the hybridization history of grapevines. Recently, haplotype-resolved T2T genome assemblies have been reported for European cultivar Yan73^13^ and Thompson Seedless^14^, although the assembly remains incomplete missing a few telomeres. Importantly, genomics has gradually shifted its paradigm from single linear reference genome to graph-based pangenome reference which can provide a comprehensive genomic variation repertoire of a species or genus^15^. Pangenome reference integrating multiple individuals from diverse genetic backgrounds has been reported in human^16^, animals^17–19^, and plants^20–23^. However, most pangenomic studies either rely on incomplete reference genomes, or ignore haplotype variations in diploid or polyploid organisms, thus inevitably missing potentially functional variations. Recently, a grapevine pangenome research was reported using haplotype genomes of 9 North American grapevine accessions^24^. However, a genus-wide pangenome reference based on high-quality genome assemblies of grapevines from all major continents is still lacking, which can serve as an essential resource for understanding the vast genomic diversity, functional genomics research, identification of lost heritability, and precision improvement of grapevines.

Here, we present the largest pangenome study of *Vitis* genus constructed from assembled and annotated haplotype-aware genomes of 72 grapevine accessions from 19 *Vitis* species, including new genome sequences for 13 *Vitis* species for the first time, as well as the first haplotype-resolved complete genome of a major winegrape *V. vinifera* cv. Chardonnay. We also reported a transcriptomic profiling of 113 *Vitis* accessions infected by downy mildew (DM) to couple with graph pangenome for genome-wide association study. Our results revealed a comprehensive genetic architecture of the *Vitis* genus and demonstrated the power of using the haplotype-resolved pangenome in combination with variome, transcriptome and phenome to identify genomic variations associated to agronomic traits, such as DM resistance and water use efficiency, thus boosting genomics-assisted breeding and genetic studies for grapevines.

## Results

### Haplotype-resolved complete genome and centromere landscape of *V. vinifera*

We first assembled a telomere-to-telomere (T2T) haplotype-resolved genome of a major winegrape (*V. vinifera*) cultivar Chardonnay as a reference for downstream variant callings. Chardonnay had an estimated genome size of 508.07 Mb and a heterozygosity rate of 1.64% based on kmer frequency analysis (**Figure S1A**). We deeply sequenced Chardonnay to generate a combination of PacBio high-fidelity (HiFi) long reads (308×), Oxford Nanopore technology (ONT) ultra-long (N50 > 100 kb) reads (117×), high-throughput chromatin conformation capture sequencing (Hi-C) reads (180×) for T2T genome assembly. From HiFi and Hi-C reads, we used hifiasm to successfully assemble two phased assemblies (hap1 and hap2). After decontamination of plastid and microbial sequences the two phased assemblies were scaffolded into pseudochromosomes using Hi-C data by Juicer^25^ and 3d-DNA^26^ yielding 19 chromosomes for each haplotype containing 3 and 0 gaps, respectively. The chromosome-level assemblies were then subjected to gap-filling using HiFi and ONT reads as well as unplaced contigs to achieve gap-free assemblies. The two final haplotype-resolved T2T genomes (VHP-T2T) sized 501.30 Mb and 503.73 Mb and each contained 19 gap-free chromosomes (Figure S1B) with all 38 telomeres (**Table S1**) with contig N50 of 26.39 Mb and 26.16 Mb, respectively (**Figure 1A and Table S1**). Both haplotypes of VHP-T2T were well aligned against a previous *V. vinifera* genome PN40024^27^ (**Figure S1E**). Extensive validations using multiple metrics such as read mapping, Hi-C interaction maps, BUSCO, LAI and QV (see methods) demonstrated the completeness and accuracy of two haploid genomes (**Table 1; Figure S1C and S1D**). All 19 centromeres with genomic locations and boundaries were clearly validated for both haplotypes of VHP-T2T by ChIP-seq using grapevine-specific antibody of CENH3 (centromere-specific histone 3) (**Figure 1A**).

**Figure 1.**
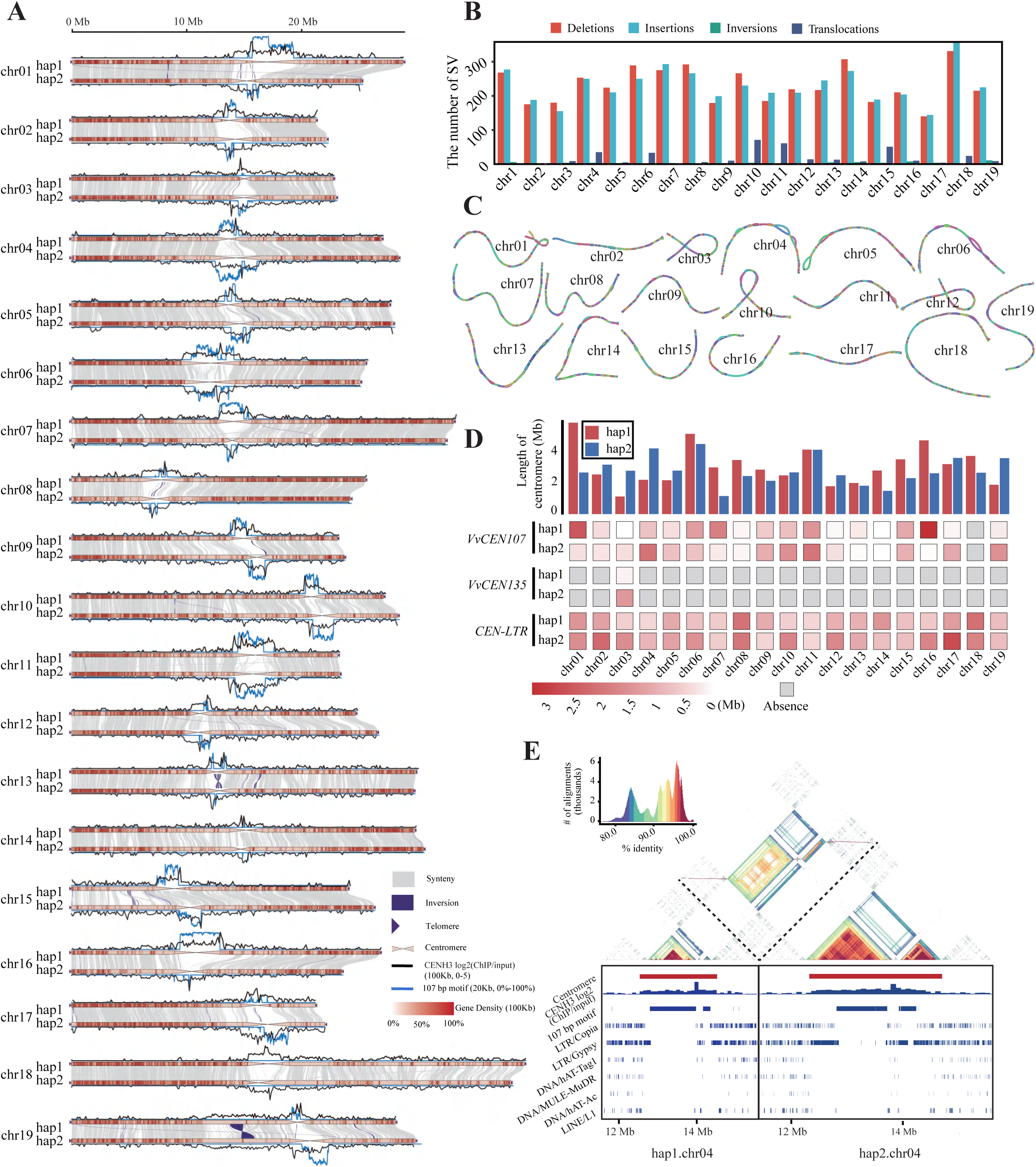
Haplotype-resolved complete genome sequence of *Vitis vinifera* cv. Chardonnay. A. Ideogram of T2T assembly for two haplotypes, on which various genomic features are visualized. For each chromosome (chr), synteny and structural variants (SVs) between hap1 and hap2, telomeres, centromeres, CENH3 log2(ChIP/input) distribution, 107 bp satellite repeat distribution, and gene density are shown. B. Bar graph summarizing the number of SVs between two haplotypes of VHP-T2T. C. Bubble plots showing the heterozygous segments between two haplotypes for 19 chromosomes. Alternating colors indicate a switch of homozygous-heterozygous status. D. Centromere length (bar graph) and motif contents (heatmap) of VHP-T2T hap1 and hap2 genomes. E. Top: a sequence identity heatmap created by StainGlass using centromere sequence of hap1 and hap2 from Chr04. Bottom: heatmaps in the two boxes shows centromere regions, CENH3 ChIP-seq signals, 107 bp motif distribution and transposon content for Chr04 centromere for VHP-T2T hap1 and hap2.

**Table 1.** Summary statistics from the haplotype-resolved T2T gap-free genome assembly and annotation of *Vitis vinifera* cv. Chardonnay.

|  | Haplotype 1 | Haplotype 2 |
| --- | --- | --- |
| **No. of contigs (gap)** | 19 (0) | 19 (0) |
| **Assembly size (Mb)** | 501.30 | 503.73 |
| **No. of centromeres** | 19 | 19 |
| **No. of telomeres** | 38 | 38 |
| **Contig N50 (Mb)** | 26.39 | 26.16 |
| **Contig N90 (Mb)** | 18.40 | 22.46 |
| **Largest contig (Mb)** | 38.76 | 37.63 |
| **BUSCO (%)** | 96.8 | 97.2 |
| **NGS mapping ratio (%)** | 99.28 | 99.27 |
| **HiFi mapping ratio (%)** | 99.91 | 99.96 |
| **ONT mapping ratio (%)** | 100.00 | 100.00 |
| **LAI (LTR Assembly Index)** | 18.92 | 17.71 |
| **QV (Quality Value)** | 72.93 | 73.36 |
| **Protein-coding genes** | 39,261 | 39,056 |
| **Repeat content (%)** | 53.56 | 53.63 |

Although the two haplotypes of VHP-T2T showed high collinearity with a sequence identity of 93.37%, we identified 9,223 structural variants (SVs) including 4,407 deletions, 4,374 insertions, 73 inversions and 369 translocations between them (**Figure 1B**), highlighting the heterozygosity nature of cultivar Chardonnay (**Figure 1C**). These variations spanned 7.8 Mb and 7.11 Mb representing 1.55% and 1.41% of hap1 and hap2, respectively. The largest difference resided in their centromere regions (**Table S2)** with a diverse repeat composition and variable centromere lengths both within and among chromosomes (**Figure 1D**), as demonstrated by the CENH3 ChIP-seq signals (**Figure 1A**). Consistent with a previous report^12^, the satellite DNA monomer of 107bp (*VvCEN107*) was the dominant repeat unit (hap1: 26.32%, hap2: 25.99%) in all *V. vinifera* centromeres. Besides, *VvCEN135* was occasionally found in a few centromeres and all centromeres were invaded by long terminal repeats (LTRs) (**Figure 1D**). For example, two haplotypes differed greatly in Chr04 centromere where the size of centromeres, *VvCEN107* copy numbers and the extent of LTR invasion were markedly distinctive (**Figure 1E**). Comparative analysis across 19 chromosomes revealed low sequence conservation among centromeres (**Figure S2**), suggesting highly variable and rapidly evolved winegrape centromeres even between two parent haplotypes. Genome annotation combining *ab initio* prediction, homology proteins and multi-tissue transcriptome data predicted 39,261 and 39,056 protein-coding genes for hap1 and hap2, respectively (**Table 1**), comparable to 37,534 in the previous report for winegrapes^12^. VHP-T2T hap1, with a higher contig N50 than hap2, was used as reference in downstream variant calling and pangenome construction.

### Genetic and phenotypic diversity of grapevines

To understand genetic and phenotypic diversity of *Vitis* genus, we collected and curated 71 diploid grapevine accessions including 25 wild grapevines and 46 cultivars, covering 19 species (**Figure 2A and Table S3**). The wild accessions included 23 East Asian and 2 North American grapevines, while the cultivars were grouped into 22 European, 19 North American, 1 East Asian and 4 Muscadine grapevines (*V. rotundifolia*) according to our phylogenetic analysis, representing a broad spectrum of grapevine genetic diversity (see below **Figure 2D and Table S3**). This collection of grapevines displayed large phenotypic variations such as leaf morphology (sizes and shapes), berry morphology, resistance to downy mildew, and stomatal density, length and conductance (**Figure 2B, 2C, S3B and Table S4**). For instance, compared to Muscadine and East Asian groups, North American and European groups had longer stomata, higher stomatal conductance and were less resistant to downy mildew, whereas Muscadine groups had higher stomatal density than other groups (**Figure S3B and Table S4**).

**Figure 2.**
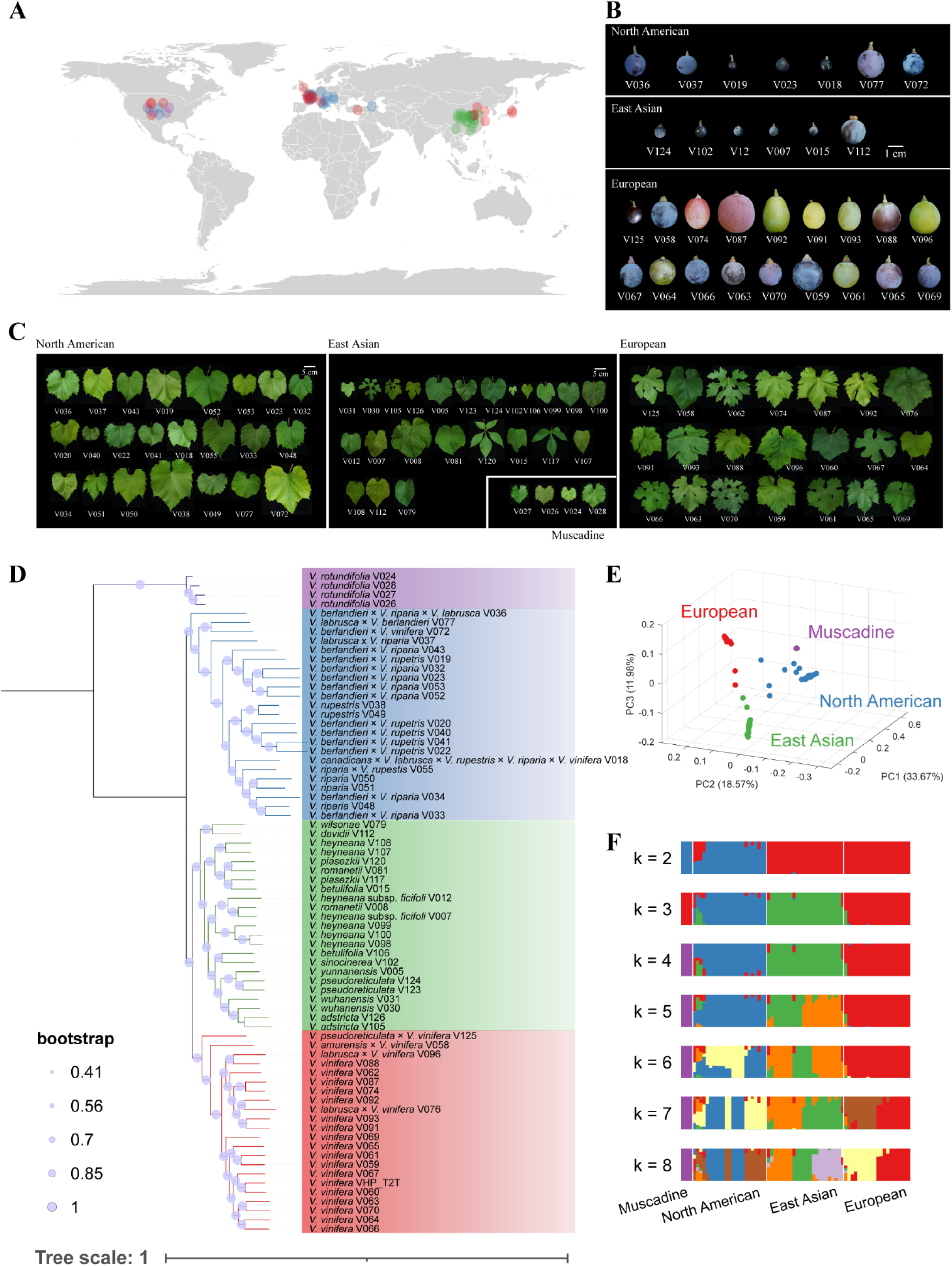
Geographic distribution and phylogenetic analysis of grapevine accessions. A. Geographic distribution of the grapevines used in this study (Purple: Muscadine, Blue: North American, Green: East Asian, Red: European originated grapevines). The map is generated by map data function in the R package ggplot2. B-C. Representative berry shape (B) and leaf morphology (C) pictures of grapevines. D. Coalescence maximum likelihood phylogenetic tree. The tree is rooted by the Muscadine clade. E. Principal component analysis of grapevine accessions. F. ADMIXTURE ancestry analysis of grapevine accessions.

We then performed population genomic sequencing of these grapevine accessions using Illumina paired-end (NGS) short reads (∼50×coverage) (**Table S5**) to detect single nucleotide polymorphisms (SNPs) and Indels using VHP-T2T hap1 as a reference genome. In total, 123 million SNPs and Indels were detected and further filtering yielded a set of 683,041 high-quality synonymous SNPs for the 71 accessions, which were used in downstream phylogenetics, principal component analysis (PCA) and biogeographic history (ADMIXTURE) investigation. Both approximately-maximum-likelihood tree (**Figure 2D**) and PCA (**Figure 2E**) showed an unambiguous separation of the 71 accessions into four clusters representing the North American, East Asian, European and the Muscadine grapevines. We further conducted ADMIXTURE analysis to understand the genetic ancestry of the 71 accessions. At k=4, all 71 grapevine accessions were divided into four distinctive ancestry groups corresponding to a single geographic group, indicating the independent origin of grapevines for the three continents as reported previously^6^ (**Figure 2F**). At k=6, the best k value determined by cross validation error (**Figure S3C**), grapevines from North America and Europe showed multiple ancestries but overall shared no ancestry among groups, suggesting a lack of genetic exchanges among them (**Figure 2F**). It also showed that the genetic diversity of North American and East Asian grapevines has not been well explored in the breeding of cultivated winegrapes (*V. vinifera*), highlighting the great potential of these resources in genetic improvement of grapevines. Meanwhile, we downloaded NGS data of 475 grapevine and outgroup accessions reported by Liang *et al.*^6^ to compile a total of 591 accessions (including 71 accessions in **Table S3** and 45 additional samples in **Table S4**) covering both wild and cultivated grapevines from the three major continents in our analysis. SNP-based phylogenetic tree of the 591 accessions suggested that our 71 grapevine accessions were a good representation of genetic diversity in the *Vitis* genus (**Figure S3A**), and thus they were used to construct the pangenome.

### Pangenome assembly and annotation of 71 grapevine accessions

For pangenome reference construction, we assembled genomes of the 71 grapevine accessions with estimated heterozygosity rates of 1.13% to 3.13% and genome sizes of 337 Mb to 483 Mb based on kmer analysis (**Table S6**). We generated PacBio HiFi (∼60×) and Hi-C (∼100×) reads (**Table S5**) for each of our 71 grapevine accessions and assembled two haploid genomes per accession with 74 to 3,057 haplotigs using hifiasm (**Table S7**), which were further anchored to chromosomes using Hi-C data. The final assemblies of 142 haplotype genomes (71 accessions x 2 haplotypes) sized from 435.8Mb to 651.2Mb (**Figure 3A**), with BUSCO scores between 93% and 97.5%, Quality Value (QV) of 34.8-71.6, and LTR Assembly Index (LAI) of 12.6-21.5 (**Figure S4A and Table S7),** suggesting high level of completeness. All haplotype-resolved genomes contained 19 chromosomes of variable sizes (**Figure S4B**) except for Muscadine grapevines with 20 chromosomes as previously reported^28^. The contig N50 of 142 haplotype genomes varied from 5.01 Mb to 26.15 Mb with a mean value of 19.36 Mb, almost comparable to the VHP-T2T assembly (**Figure 3B**). Genome annotation of the 142 haplotypes integrating *ab initio* prediction, homolog and transcriptome evidence predicted 31,761 to 36,639 (mean: 35,185) protein-coding genes for Muscadine, 35,403 to 42,168 (mean: 39,905) for European, 39,151 to 43,318 (mean: 40,895) for East Asian and 38,586 to 51,051 (mean: 41,177) for North American accessions (**Table S7**). Overall, two haploid genomes of all accessions had comparable genome sizes, NG50, numbers of annotated genes and repeat contents with only minor exceptions (**Figure 3A**). For example, repetitive elements occupied 36.77% to 52.73% of the genomes (**Figure 3A and Table S7**) and genome sizes were positively correlated with the LTR transposon occupancy (R^2^= 0.26-0.68) with the exception of *V. rotundifolia* (R^2^=0.12) accessions (**Figure 3C**). In addition, five haplotypes of *V. rotundifolia* accessions had unusually twice as much DNA transposon presence than the three groups of *Euvitis* grapevines (**Figure 3C and Figure S4C**). The distinctive repeat composition in Muscadine with *Euvitis* grapevines was overall in line with their phylogenetic relationships (**Figure 2D**).

**Figure 3.**
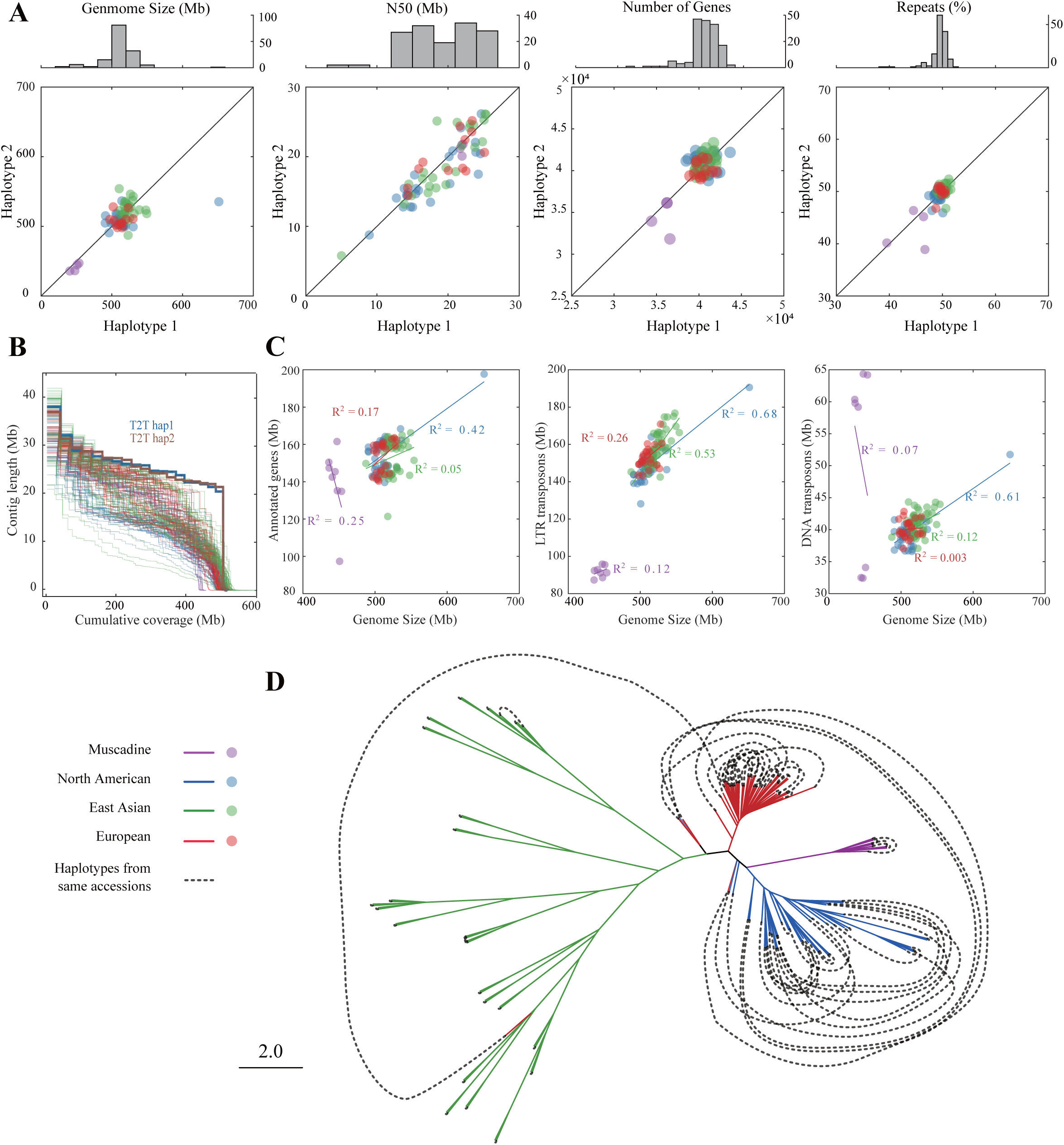
Assembly and annotation statistics, and coalescence phylogenetics of haplotype-resolved grapevine genomes. A. Genome size, NG50, annotated gene number, repetitive sequences, of two haplotypes of the same accessions. The x=y line is drawn in black. B. Assembly continuity plot of two T2T genomes and 142 chromosome-scale haplotype-resolved genomes. C. Correlation of genome size and sequence occupied by DNA transposons, LTR transposons, and genes each with a fitted line and correlation coefficient score (R^2^). D. The coalescent tree from 19 chromosome-based trees constructed with SNPs from the pangenome graph. Dotted lines connect the two haplotypes of the same accessions.

### Haplotype-aware phylogenomics reveals the hybridization history of grapevines

Haplotype-resolved genome assemblies can provide unique and unprecedented insights into the hybridization history of species even without trio information. With 144 haplotype grapevine genomes (VHP-T2T included), we constructed a whole-genome coalescent phylogenetic tree to investigate the hybridization history of these grapevine accessions. The Pangenome Graph Builder (PGGB)^29^ was first used to generate a pangenome graph of 144 haplotypes. Using SNPs from the deconstructed graph with VHP-T2T hap1 as reference, we built phylogenetic trees for each chromosome and then used ASTRAL^30^ to generate a coalescence tree from 19 chromosome-based trees (**Figure 3D and Table S8**). The coalescence tree showed a clear separation of haplotypes belonging to each of three continents and Muscadine. There was extensive intra-group hybridization of haplotypes for North American and European but not East Asian accessions. In addition, inter-group hybridization across different continents was minimal, especially between East Asian accessions and the other two continents. These results are consistent with the fact that East Asian accessions are rarely used in breeding and thus most of the samples we collected were from wild sources, while other samples were mainly from cultivars (**Table S3**). Though we cannot rule out the possibility that incompatibility exists, complete failure of hybridization among species in *Vitis* genus has not been reported to the best of our knowledge and it is generally accepted by the *Vitis* research field that *Vitis* species are inter-fertile^5,6^. Thus, genome information of these newly released wild accessions represents a resource of genetic pool for grapevine improvement.

While the diploid phylogenetic tree (**Figure 2D**) and haploid tree were mostly concordant with each other, there were a few exceptions (**Table S8**). For example, a *V. pseudoreticulata* ×*V. vinifera* hybrid V125 was grouped to the European clade in the diploid tree, whereas in haploid tree, its two haplotypes were placed in European and East Asian clade, respectively. Therefore, the results demonstrated the value of haplotype-based whole-genome phylogeny for correctly tracing back the grapevine hybridization history.

### Pangenome analysis of grapevines

With the 144 fully annotated haplotype *Vitis* genomes, we performed pangenome analysis on the 72 grapevine accessions using previously reported methods^20^, classifying all genes into a total of 64,517 gene families from 144 genomes (**Table S9**). The number of gene families increased rapidly as additional genomes were included and approached a plateau with n=125 genomes (**Figure 4A**), indicating a closed pangenome with the 72 accessions. Of the total gene sets, 8,223 families (12.75%) were present in all 144 genomes and defined as core genes, 9,252 families (14.34%) were present in 130 to 143 genomes and defined as softcore genes, 45,988 families (71.28%) were present in 2 to 129 accessions and defined as dispensable genes, 1,054 families (1.63%) were present in only one accession and defined as private genes (**Figure 4B**). Although dispensable and private gene families constituted a larger proportion (72.91%) of the total gene sets in the 72 accessions (**Figure 4B and 4C**), they only accounted for an average of 29.82% of the genes in individual accessions (**Figure 4D**). European grapevines contained significantly larger fraction of softcore and smaller fraction of private genes than North American and East Asian grapevines (**Figure S5A**), indicative of gene loss and reduced gene diversity in cultivated grapes. Core genes having lower nucleotide diversity (π) (**Figure S5B**) and dN/dS ratio indicated they were more functionally conserved than dispensable genes (**Figure S5C**). While core and softcore genes were enriched in house-keeping processes such as nucleologenesis, cell differentiation and guard cell development (**Figure S5D and S5E**), the dispensable and private genes were enriched for biosynthesis of secondary metabolites and chemotaxis (**Figure S5F and S5G**). Grapevine downy mildew is the most destructive disease in the global viticulture, caused by the oomycete *Plasmopara viticola* specifically penetrating through stomata, resulting in yield loss and reduction of grape quality. Nucleotide-binding site leucine-rich repeat (NLR) genes are key players in plant immunity against pathogens and diversified among individual accessions of plants^31^. We annotated grapevine NLR genes using NLR-annotator^32^and performed pan-NLRome analysis from the 72 grapevine accessions showing variable downy mildew resistance. In total, we discovered 104,046 NLR genes (**Table S10**) categorized into eight subfamilies (**Figure 4E**) across 144 haplotypes (**Figure 4F**), including 24,236 core, 49,784 softcore, 29,953 dispensable and 73 private NLR genes (**Figure S5H, S5I and Table S10**). These grapevine NLR genes mostly resided in gene clusters and many were located near subtelomere regions (**Figure 4E**). East Asian grapevines were overall more DM-resistant (**Figure 4G**) and harbored more NLR genes than European accessions (**Figure 4H**), especially for the subfamilies encoding coiled-coil NB-ARC domains (**Figure 4F**). These results suggested that wild grapevines from East Asian contained a larger repertoire of NLR genes than cultivated European grapes, altogether probably contributing to their differential DM resistance.

**Figure 4.**
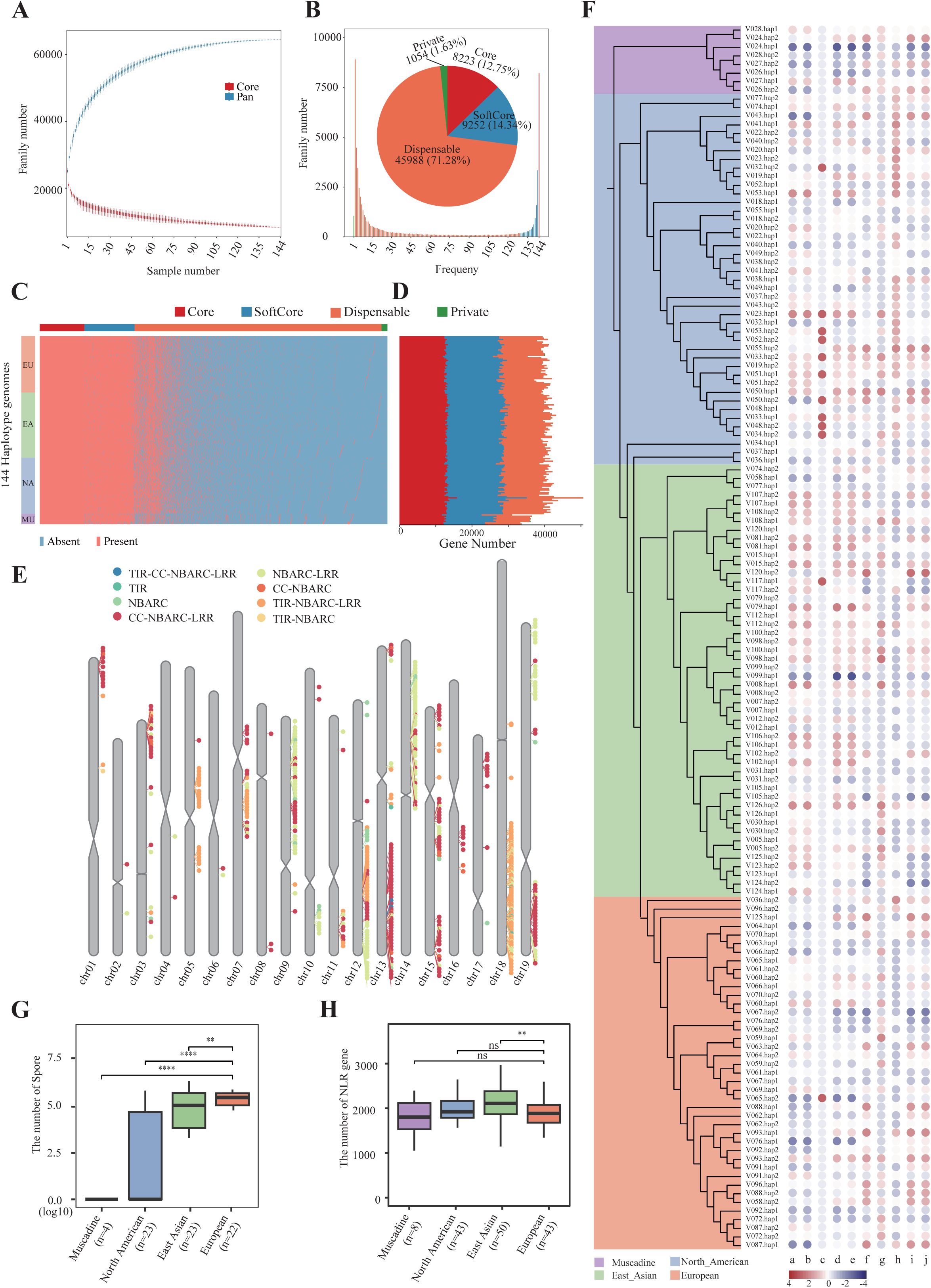
Pangenome analysis of grapevines. A. Variation of gene families in the pan- and core-genome with additional grapevine genomes. B. Compositions of the grapevine pangenome and individual haploid genomes. The histogram shows the number of gene families in the 144 haploid genomes with different frequencies where colors of histograms correspond to core, softcore, dispensable, and private genes. The pie chart shows the proportion of the gene family marked by each composition. C. Presence and absence variation of gene families in 144 haplotypes. D. Distribution of core, softcore, dispensable and private genes in 144 haplotypes. E. Chromosomal distribution of NLR genes in VHP-T2T hap1 genome. F. Distribution of NLR genes shown in a heatmap of column z-scores normalized from the number of genes. On the left, the haplotype-resolved phylogenetic tree is shown. a-j: the number of CC.NBARC, CC.NBARC.LRR, CC.TIRNBARC.LRR, NBARC, NBARC.LRR, TIR, TIR.CC.NBARC.LRR, TIR.LRR, TIRNBARC and TIR.NBARC.LRR. G. Disease index of downy mildew as shown by the number of spores after *P. viticola* (*Pv*) infection. \*\**P* < 0.01, \*\*\**P* < 0.001, \*\*\*\**P* < 0.0001, Student’s t test. H. The number of NLR genes in different grapevine groups. \*\**P* < 0.01, Student’s t test. ns, not significant.

### Extensive structural variations present in wild and cultivated grapevines

Large-scale genomic variants such as SVs are major drivers of plant evolution and domestication. We detected SVs in 67 *Euvitis* accessions (Muscadine accessions were omitted due to different numbers of chromosomes) by aligning the 135 high-quality haplotype-resolved genome assemblies (134 haplotypes + VHP-T2T hap2) against VHP-T2T hap1 as the reference using minimap, followed by SV detection using *Syri*^33^. A total of 132,518 non-redundant SVs (54,784 insertions, 62,652 deletions, 3,641 inversions, and 11,441 translocations) of 51 bp to 7,473,130 bp were detected (**Figure 5A and S6**). North American and East Asian accessions had more SVs detected than European ones (**Figure 5B**), probably due to using a reference genome of European origin (*V. vinifera*). With 135 haplotype genomes, the number of pan-SVs have not reached a plateau indicating that with larger sample set more SVs could be detected (**Figure 5C**). The majority of insertions and deletions (90% < 1,053 bp) were shorter than the inversions and translocations (90% < 313 kb), with some inversions longer than 1 Mb (**Figure 5D**). The inversions and translocations were mainly distributed in and near centromeres (**Figure S7A and S7B**), whereas deletions and insertions lacked genomic hotspots (**Figure 5A**). The large SVs (>1Mb) detected by pairwise whole-genome alignments were validated by mapping sample Hi-C reads against VHP-T2T (**Figure S7C-S7F**). For example, one 3.4Mb inversion event on Chr7 for *V. heyneana* hap1 was visible on Hi-C interaction map showing abnormal chromatin interaction signals of a typical inversion (**Figure 5E**). Although SVs were mostly located at the intergenic regions and TEs (**Figure 5F**), 67.5% were found in 3-kb upstream, downstream, and genic regions of 19,244 protein-coding genes, thus potentially disrupting their functions. Gene ontology enrichment of these genes suggested that SVs may have well contributed to the grapevine evolution in terms of leaf morphology, detection of biotic stimulus and pathogen recognition (**Figure 5G**). Particularly, our SV collection detected previously characterized^34,35^ molecular markers for downy mildew resistance routinely used in grapevine breeding (**Figure 5H, 5I, S8 and S9**). One such SV on *Rpv3* locus (UDV305)^35^ was located on the promoter of *VvSEC14* (cytosolic factor family protein) homologous to Arabidopsis *SEC14* involved in stress response^36^. This 112bp deletion was present in mostly European accessions showing increased *VvSEC14* expression and disease susceptibility (**Figure 5H and 5I**). We have also identified two additional SVs, a 59bp deletion on promoter region and 238bp insertion in 3’UTR of *VvSEC14,* mostly in North American accessions with reduced *VvSEC14* expression and disease resistance (**Figure 5I**).

**Figure 5.**
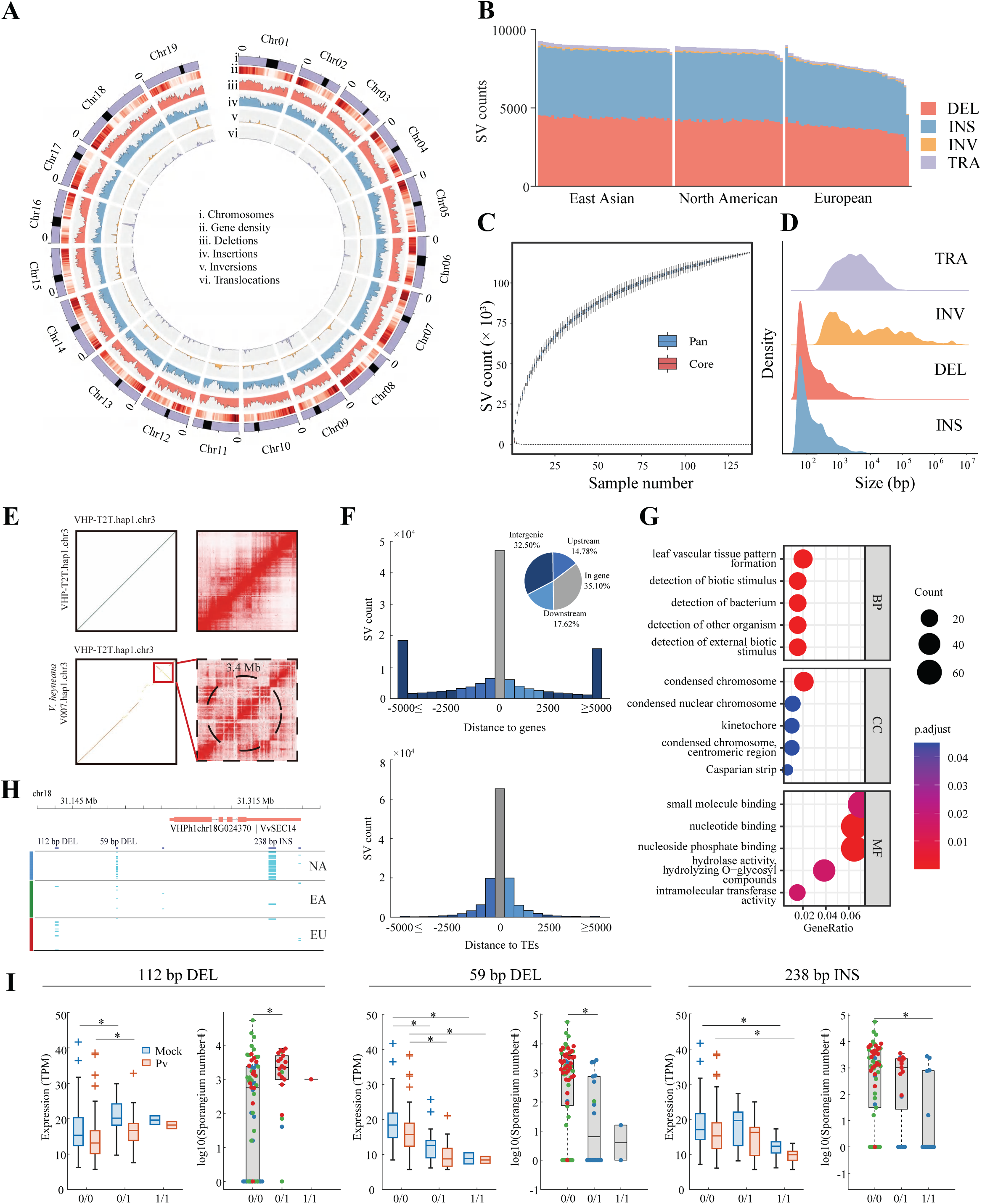
Structural variant landscape and pan-SV analysis in grapevines. A. Distribution of SVs in VHP-T2T genome. i) chromosomes (centromeres shown in black), ii) gene density, iii) deletions, iv) insertions, v) inversions, vi) translocations. B. SV number distribution in East Asian, North American, and European accessions. C. Variation of SV count in the pan- and core-SV along with additional grapevine genomes. D. SV size distribution. TRA: translocation. INV: inversion. DEL: deletion. INS: insertion. E. Example of a 3.4Mb inversion event on Chr3 of V007 sample (bottom) validated by Hi-C data, as opposed to a reference control (top). Left, dot plot showing genome alignment to VHP-T2T. Right: Hi-C contact heatmap showing Hi-C reads mapping to the VHP-T2T genome. F. SV distribution with respect to the genes (top) and transposable elements (TE, bottom). G. Enriched GO (gene ontology) terms in genes affected by SVs. BP: biological process. CC: cellular component. MF: molecular function. H. Gene model for *VvSEC14* with three SVs. The haplotypes that have the mutations are shown in cyan. NA: North American. EA: East Asian. EU: European. I. Expression levels of *VvSEC14* gene and the number of spores collected when infected with *P. viticola* (*Pv*) in accessions of different SV genotypes: no SV (0/0), with heterozygous SV (0/1), and homozygous SV (1/1). Purple: Muscadine, Blue: North American, Red: European accessions. \**P* < 0.05, Wilcoxon Rank Sum test.

### Graph-pangenome enabled discovery of SV-eQTLs for downy mildew resistance

SVs have major impacts on agronomically important traits by disrupting gene coding sequence or gene expression. However, population-level SV genotyping is challenging in plants, impeding SV–phenotype associations. Graph-based pangenomes are capable of storing both reference and alternative allele sequences while retaining the coordinate systems of the linear reference genome, which facilitates mapping of short reads from SV regions and thus SV genotyping. Several studies reported SV-genotyping by mapping low coverage NGS reads to graph-based pangenome for SV genotyping^22,37^. We constructed a grapevine graph-based genome by integrating the linear reference genome sequence of *V. vinifera* VHP-T2T and the 132,518 non-redundant SVs identified from the 67 *Euvitis* genomes using *vg* pipeline (**Figure 6A**). The resulting graph pangenome had 535,122,987 bases containing 16,830,980 nodes and 16,946,222 edges. We then genotyped these SVs in 113 accessions (**Table S4**) by mapping their NGS data (50x) against the pangenome graph using *Giraffe* and *vg*^38^. With the genotyped SVs, we performed expression quantitative trait loci (eQTL) analysis by incorporating transcriptome data of 113 accessions infected by *P. viticola* at 1 dpi (days post inoculation) (**Table S4**). The expressed genes (TPM >1) and SV genotypes were used in eQTL analysis as previously described^39^, which identified 79 significant association signals (*P* < 2.14e-5) (**Figure 6B and Table S11**).

**Figure 6.**
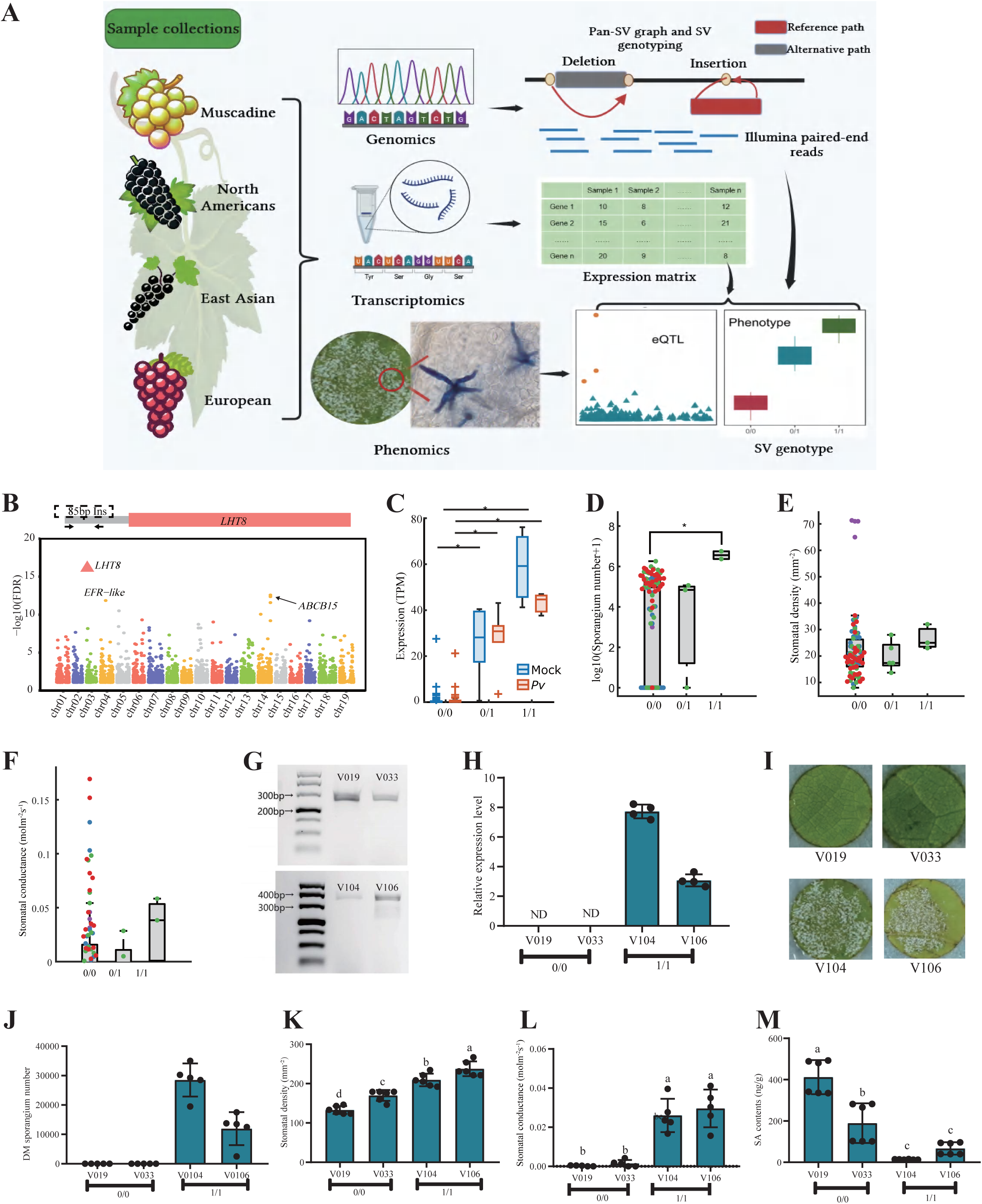
Graph pangenome and multi-omics identified SV-eQTLs for downy mildew resistance. A. Schematic flowchart of grape pangenome construction and SV-eQTL analysis. B. Manhattan plot of SV-eQTL analysis for downy mildew (DM) resistance. The gene model of *LHT8* gene is shown with the primers (black arrows) used for PCR validation of the 85bp insertion significantly linked to DM resistance. Two other significant linked SV-eQTLs (*EFR-like* and *ABCB15*) were also marked with arrows. Dotted horizontal line indicates significance threshold. C-F. Expression levels (C), sporangium numbers after *P. viticola* (*Pv*) infection (D), stomatal density (E), and stomatal conductance at 10 pm (F) in accessions of different SV genotypes. \**P* < 0.05, Wilcoxon Rank Sum test. G. PCR validation of the 85bp insertion in accessions with (bottom) or without SV (top). H-I. Relative expression levels (H), representative leaf images showing DM sporangium (I) in accessions infected by *P. viticola* (*Pv*) at 7 days post inoculation. J-M. Sporangium number (J), stomatal density (K), stomatal conductance at 10 PM (L), and salicylic acid contents (M) in the accessions with homozygous SV or without SV. Different letters indicate statistical significance (*P* < 0.05, ANOVA with Tukey’s test); ND, not detected. 0/0, no SV; 0/1, with heterozygous SV; 1/1, homozygous SV.

The most significant signal was located at Chr3 where a 85 bp insertion event was strongly associated with expression level changes for a gene encoding a leucine histidine transporter homologous to *Arabidopsis thaliana LHT8* (*VvLHT8* hereafter) which was reported to regulate plant defense response against bacterial and fungal disease in a salicylic acid (SA)-dependent manner^40^. The *VvLHT8* expression was higher in the accessions with 85 bp insertion than these without (**Figure 6C**) and those SV-homozygous accessions were more susceptible to DM (**Figure 6D**) and had different stomatal traits including higher stomatal density and stomatal conductance at night (**Figure 6E-6F**). The results were validated with grapevines either without the SV or being SV-homozygous (**Figure 6G-6L**). We also found that the SV-homozygous accessions showed lower SA content in the leaves (**Figure 6M**). Stomata are key players in not only water use efficiency but also warding off pathogen invasion, and SA is an important phytohormone regulating stomatal movement and plant immunity^41–43^. These results suggested that *VvLHT8* could regulate DM resistance and water use efficiency in a stomate- and SA-dependent manner. In addition to *VvLHT8*, expression levels of several other genes from the eQTL analysis were also found significantly related to DM resistance and stomatal traits (**Figure S10**), and many others can be identified through further analysis of the phenotyping data of the 113 grapevines provided in **Table S4 and S11**. These genes are thus strong genetic targets for improving DM resistance and water use efficiency of grapevines.

## Discussions

Grapevine is a model species for studying plant domestication and evolution^1,7^ and is also an important cash crop with over seven million hectares planted across the globe, contributing over hundreds of billion dollars to the world economy^44^. Surprisingly, grapevine genomics, once in frontiers, have been progressing much slower than other crops, largely because of their high genome heterozygosity caused by extensive hybridization. Despite several published reference genomes of grapevines^8–11,13,14,24,27^, complete and haplotype-aware reference genomes remain lacking to date. Using sequencing data generated from the latest PacBio HiFi, ONT ultra-long and Hi-C technologies, we assembled the first haplotype-resolved complete genome of *V. vinifera* cv. Chardonnay, the most influential grapevine for making white wine. It allowed us to gain insight into the substantial genomic variations between two haplotype genomes including CENH3-ChIP validated centromere regions, revealing unprecedented landscape of high-copy tandem repeats within haplotype-aware grape centromeres. This opens the path to dissect the function and evolution of centromeres in grapevine genomes which have long served as the pillars for plant evolutionary genomics studies. We further explored haplotype landscape of 71 global grapevine accessions (25 wild + 46 cultivars) representing the broad genetic diversity of the genus by producing high-quality haplotype-resolved genomes of them to construct a pangenome and trace the complex hybridization of grapevines. To the best of our knowledge, genomes of 60 *Vitis* accessions in our sampling are newly released including new genomes for 13 *Vitis* species, while genomes of the other 12 *Vitis* accessions have recently been reported ^9,13,14,45–50^ but further improved in this study (**Table S3**). This is so far the largest pangenome study integrating genus-wide haplotype-aware genomes reported for any plants. The genomes of 47 cultivars will directly facilitate the research and breeding of grapevines and thus benefit the grape industry.

Human domestication of grapevines dates back to ∼11,000 BC and is primarily based on a single species *V. vinifera*^1^. This has led to a rather narrow genetic diversity among grapevine cultivars making them susceptible to various biotic and abiotic stresses. Therefore, shifting the attention to wild grapevine species for genetic and genomic research is critical to improving grape cultivars. In this work, we released haplotype-resolved reference genomes of 25 new wild grapevines, which are crucial germplasms for improving the cultivars in many traits, particularly resistance to biotic and abiotic stresses. The current grapevine breeding is heavily dependent on the crossing between European and North American grapevines, which was clearly supported by our haplotype-based coalescent phylogenetic analysis (**Figure 3D**). The same analysis also highlighted that Asian grapevines were poorly explored in breeding and represent a valuable genomic resource for grapevine improvement. We expect these high-quality grapevine genomes to empower future research endeavors towards answering key questions about grapevine biology and evolution, such as the mechanisms behind heterosis, sex locus and centromere evolution etc.

Grapevine genomics has been relying on a single linear reference genome of European grapevine, which introduces reference bias owing to its narrow representation of grapevine diversity. From the assembled and annotated *Vitis* genomes, we built a graph-based pangenome reference for grapevine functional genomics research and molecular breeding, a hallmark paradigm shift from current approach. By including a published genome of the wild European grapevine *V. vinifera* ssp. *sylvestris*, our pangenome covers the most comprehensive genome information of *Vitis* genus to date (**Figure 4**), which is supported by the saturated number of identified core and dispensable gene families in our grapevine collection. We also extensively characterized SVs in each of the grapevines, obtaining the largest catalog of structural variations for *Vitis* spp. The high-quality genomes, variomes, phenomes, transcriptomes from global grapevine accessions generated in this study will make our grapevine collection a useful material pool for both functional genomics studies and breeding of grapevines. Identification of genetic markers, particularly specific gene sequences, is key to genomics-assisted and gene-editing based crop improvement. To show the power of pangenome in identification of trait-associated genetic markers, we focused on critical agronomic traits, DM resistance and stomatal traits closely relevant to water use efficiency, and identified 79 SVs strongly linked to these traits using SV-eQTL analysis (**Figure 6**, **S10 and Table S4)**. By analyzing SVs in the known DM-resistant QTLs, we identified *VvSEC14* in the *Rpv3* locus^34,35^ negatively regulating DM resistance (**Figure 5I**), thus representing a potential target to improve usage of the locus. Further functional studies of the genes affected by the SVs will facilitate precision improvement of grapevines. There have been several large-scale genomic resequencing and phenotyping projects of grapevines^6^. Population-scale variant detection and genotyping by mapping resequencing reads to our pangenome followed by GWAS is a cost-saving and unbiased approach that will accelerate the identification of trait-associated genetic markers. Overall, the current study released the first haplotype-resolved complete genome and the largest pangenome of grapevines, revealed the genetic architecture of the *Vitis* genus and demonstrated the power of employing the pangenome to uncover genomic variants linked to important grapevine traits such as DM resistance and water use efficiency, thus paving the way for future research and marker-assisted improvement of grapevines towards the goal of generating a super grapevine for human beings.

## Online Methods and Materials

### Plant resources and sample collection

The *Vitis* accessions used in this study were grown in the vineyards at Zhengzhou Fruit Research Institute, Chinese Academy of Sciences, and Institute of Advanced Agricultural Sciences, Peking University, China. Standard management procedures, such as cultivation, watering, fertilization, pruning, and disease control, were applied to all plants.

### DNA and RNA isolation

Genomic DNA used for NGS short-read sequencing was isolated from grape plants using the DNeasy Plant Mini Kit (QIAGEN). The Agilent 4200 Bioanalyzer (Agilent Technologies, Palo Alto, California) was used to evaluate the DNA’s integrity. High molecular weight DNA used for long-read sequencing was isolated from fresh leaves of grapevines using CTAB (cetyltrimethylammonium bromide) method as described previously^51^. Genomic DNA quality was assessed using 1% agarose electrophoresis to determine whether the samples were contaminated and degraded. The purity of the samples was detected using a NanoPhotometer® spectrophotometer (IMPLEN, CA, USA) and DNA sample concentration was determined by the Qubit® 2.0 Flurometer (Life Technologies, CA, USA). Total RNA was isolated from grapevine leaves using TRIzol reagent (Thermo Fisher, USA) following manufacturer protocol. The concentration of extracted nucleic acid was detected with Nanodrop2000 (Thermo Fisher, USA). For RNA quality examination, the purity of the samples was determined by NanoPhotometer ® (IMPLEN, CA, USA), while the concentration and integrity of RNA samples were detected by Agilent 2100 RNA nano 6000 assay kit (Agilent Technologies, CA, USA).

### Genome sequencing

For NGS sequencing library, g-Tubes (Covaris) were used to shear 15 µg of genomic DNA and AMPure PB magnetic beads were used to concentrate the DNA. Library construction was conducted using a 1µg gDNA template according to the steps sepecified in the TruSeq DNA Sample Preparation Guide (Illumina, 15026486 Rev.C). HiSeq X Ten sequencing platform (Biomarker Inc, Qingdao) was used to execute the paired-end sequencing program (PE150) to produce 150 bp sequence reads. For standard Hi-C library preparation, young leaves were cross-linked by 40 ml of 2% formaldehyde solution at ambient temperature in a vacuum for 15 min, following a standard Hi-C protocol^52^. The constructed Hi-C sequencing library was first subjected to a test run of sequencing to evaluate valid interaction read pairs using HiC-Pro (v3.1.0)^53^ before high coverage sequencing by Illumina NovaSeq 6000 to yield 50∼126 Gb (126∼334×genome coverage) paired-end reads. To generate PacBio HiFi data, a total of 15 µg purified HMW genomic DNA were used to construct a standard PacBio SMRTbell library using PacBio SMRT Express Template Prep Kit 2.0 (Pacific Biosciences, CA). The CCS (circular consensus sequencing) was performed using a PacBio Sequel IIe instrument at Biomarker Technologies Corporation (QingDao, China) to generate 24∼156 Gb (58∼308×genome coverage) HiFi reads with read length N50 of 13.4∼23.5 kb. To generate Oxford Nanopore ultra-long reads, the long DNA fragments were size-selected and processed following the Ligation Sequencing SQK-LSK109 Kit (Oxford Nanopore Technologies, Oxford, UK) protocol. The final DNA library was sequenced using the PromethION sequencer (Oxford Nanopore Technologies, Oxford, UK) by Single-Molecule Sequencing Platform at Annoroad Inc. (Beijing, China).

### Genome assembly

For genome survey to estimate genome sizes and heterozygosity of 72 grapevine accessions, Illumina paired-end reads were used in kmer frequency analysis with Jellyfish v2.3.0^54^ (k-mer=21) and Genomescope v1.0^55^. To assemble the haplotype-resolved T2T genome for *V. vinifera*, high coverage PacBio HiFi (308×) and ONT ultra-long (117×) reads were used to produce an initial assembly of two haplotypes using *hifiasm* v0.19.8^56^ ‘--hom-cov --h1 --h2 --ul’ by incorporating Hi-C reads. The initial assemblies of two haplotypes were then scaffolded using Hi-C reads by Juicer^57^ and 3d-DNA^58^ pipeline. Briefly, HiC-Pro^53^ was used to extract valid read pairs after the Hi-C data was aligned to the draft genomes using Bowtie2 v2.2.3^59^. The alignment was then used to calculate the Hi-C interaction matrix by Juicer and the assemblies were scaffolded using 3d-DNA. On the basis of signals from Hi-C interaction heatmaps, manual adjustments were applied to the scaffolded pseudochromosomes using Juicebox v1.11.08^57^. ONT ultra-long reads were used in gap-filling of the chromosome-scale assemblies. The ONT reads first underwent filtering by fitlong v0.2.1 (https://github.com/rrwick/Filtlong) with parameters --min_length 80000 --min_mean_q 9. Then, consistency correction was performed on the filtered reads using NECAT version 0.0.1. Using the corrected ONT and HiFi reads, TGS-gapcloser v1.2.1^60^ with the parameter --tgstype ont --ne, was used to close gaps in the scaffolded genomes. And contigs not incorporated into the scaffolds were aligned to the chromosomes using minimap2 ‘-x asm5’. Sequences intended for gap filling must satisfy two criteria: 1) the contig should span the entire gap region, 2) the sequences on both sides of the gap, 100kb away from it, should exhibit over 90% sequence identity with the contig, and the alignment length should exceed 100kb. Subsequently, the filled positions were examined in IGV (Integrative Genomics Viewer) for validation of correctness. Genome assemblies of other 71 grapevine accessions were generated from HiFi and Hi-C reads using similar procedure as the T2T genome assembly described above except that no ONT and HiFi reads were used for gap-filling. After the initial haplotig-level assembly, the haplotigs were anchored to chromosomes using Hi-C reads of each accession by Juicer v1.6^57^, 3D-DNA v201008^58^, and Juicebox v1.11.08^61^. To remove plastid sequences from all genome assemblies produced in this study, the genome contigs were aligned to *V. vinifera* mitochondrial and chloroplast genome sequences (GenBank accessions FM179380.1 and DQ424856.1, respectively) with minimap2 v2.24^62^ with parameter ‘-x asm5’. Contigs with base coverage of at least 50% identical with a plastid sequence were eliminated from the final assemblies. Other common contamination sequences (e.g. microbial DNAs) were further eliminated by aligning the contigs to all of the RefSeq bacterial genomes from NCBI with blast-2.13.0+ with parameters ‘-task megablast’.

### Genome polish and validation

To polish genome assemblies PacBio HiFi and ONT ultra-long reads were aligned to the assemblies using Winnowmap^63^, retaining only the primary alignments. Small indels (< 50bp) were called from the alignment results of HiFi using DeepVariant^64^. Structural variations (> 50bp) were identified from the alignment results of Hifi and ONT (for VHP-T2T) using Sniffles. Merfin was then applied to evaluate the variant calls and BCFtools^65^ was used to correct the *bona fide* variants in the genome assemblies. To validate the VHP-T2T assemblies, we remapped the raw sequencing reads to the genome assemblies, and examined the mapping results for abnormal coverage of reads genomewide. Briefly, a total of 156.9 Gb of HiFi reads were aligned to the haplotype-resolved T2T genome using Winnowmap with the parameters ‘-W -ax map-pb’. For subsequent read depth analysis, only primary alignments with MAPQ value >30 were considered. The read depth for each base was then calculated using samtools (v.1.6)^66^ with the parameter ‘bedcov’. Afterwards, average depths were computed for all 1 kb bins across the genome. Bins with depths < 150 and > 600 were designated as low-coverage and high-coverage regions, respectively. Additionally, we aligned 29.9 Gb of high-quality ONT ultra-long reads to the genome using the same approach. Bins with depths < 50 and >200 were respectively defined as low-coverage and high-coverage regions, respectively. We found the low-coverage regions (4.3% of total genome) were generally associated with large segments of satellite arrays such as centromeric regions, and high-coverage regions (0.15% of total genome) were enriched with plastid DNA insertion regions. Nonetheless, we found a generally normal coverage (expected from sequencing depths) of mapped reads across all genomic regions, suggesting correct assembly of genomes even at complex regions. Then, we also looked for any large and obvious assembly errors by checking Hi-C interaction maps and found none. Finally, several standard quality control metrics were applied to evaluate the accuracy and completeness of final genome assemblies as follows. BUSCO v5.4.2^67^ was used to evaluate the genic region completeness. Using LTR retriever v2.9.0, the completeness of intergenic regions was assessed by LAI^68^. Lastly, k-mer based the quality of the assemblies was assessed by Meryl v1.3 and Merqury v1.3^69^ using default parameters.

### Annotation of pangenome

Genome repeats were annotated using RepeatModeler (v1.0.11)^70^, followed by genome soft-masking through RepeatMasker (v4.1.2.p1)^71^. The prediction of the gene model utilized an integrated pipeline that included the following data: *ab initio* predictions, homologous protein comparisons, RNA-seq, and Iso-seq evidence. In the *ab initio* prediction phase, we trained the GeneMark-ET model using BRAKER2 (v.2.1.6)^72^ and further refined it by training the semi-HMM model SNAP^73^ via MAKER (v3.01.03)^74^. The plant homologous protein sequences used in this work were non-redundant, human-curated, and retrieved from the UniProt Swiss-Prot database (https://www.uniprot.org/downloads) to ensure their uniqueness. To further increase the dataset’s comprehensiveness, we also included published peptide sequences from *Vitis* as mentioned in Massonnet et al.^75^, Jaillon et al.^7^, and Minio et al.^76^. We refined the homologous protein set using cd-hit (version 4.8.1)^77^ with default parameters to reduce potential redundancy. Transcriptome evidence was included in genome annotation integrating the leaf transcriptome of the 71 grapevine accessions (**Table S3**) and multi-tissue transcriptomic data from Vitis species downloaded from NCBI (**Table S12**). The public transcriptome data were categorized into three groups by population of three origins: Europe, North America, Asia. Subsequently, each group of transcriptome data was utilized for gene annotation for the accessions of the corresponding population. T2T genomes were annotated using transcriptomic data from multiple tissues of Chardonnay cultivar. Trinity (v2.8.4)^78^ was used for *de novo* transcriptome assembly^78^. The final gene model was predicted by MAKER pipeline with protein-coding genes having an AED < 0.5.

### Centromere and telomere identification

Telomere sequences were detected using Tandem Repeat Finder (TRF) v4.09.1^79^ with the following parameters: ‘2 7 7 80 10 50 500 -d -h ‘. The generated ‘.dat file’ was converted into a GFF3 file format by using TRF2GFF (https://github.com/Adamtaranto/TRF2GFF), making it easier to recognize the seven-base telomeric repeats. For *in silico* centromere identification, tandem repeat annotation was carried out using TRASH^80^, with the parameter configuration as ‘2 7 7 80 10 50 500 -f -d -m’. The relative positions within the centromere were aligned with blast-2.13.0+ concurrently by utilizing the tandem repeat sequence units of the centromeres identified by Huang et al.^81^. The colorful identity heatmaps of genomic sequence were generated using StainGlass^82^.

### CENH3 ChIP-seq and data analysis

ChIP was performed according to a described previously method, standardized with anti-grape-CENH3 antibodies. An antigen with the peptide sequence ‘RTKHPAVRKTKALPKK’ corresponding to the N terminus of grape CENH3 was used to produce the antibody. For ChIP experiment, seedlings were fixed with 1% formaldehyde solution in MS buffer (10 mM potassium phosphate, pH 7.0; 50 mM NaCl) at room temperature for 15 min in a vacuum. After fixation, the seedlings were incubated at room temperature for 5 min under vacuum with 0.15 M glycine. Approximately 1 g fixed tissue was homogenized with liquid nitrogen and purifying nuclei, and resuspended in 1 ml of cell lysis buffer & incubate for 10 min on ice & spin at 1500 rpm (RC-3B, 600 x g) for 5 min (Cell lysis buffer: 10 mM Tris, 10 mM NaCl, 0.2 % NP-40 [pH 8.0], 1X protease inhibitors). The cell lysis was further resuspended in 1 ml of nuclei lysis buffer for 10 min on ice (Nuclei lysis buffer: 50 mM Tris, 10 mM EDTA, 1% SDS, 1X protease inhibitors) to isolate nuclei. The resuspended chromatin solution was sonicated five times for 15 s each at ∼10% power (setting 2.5 on the sonicator, Sanyo Soniprep 150). The volume of the chromatin sample was measured and then ChIP dilution buffer was added to 1ml of chromatin with 2.5 µg of anti-H3K4me3 and incubated for 12 h at 4°C. 50 µl protein A/G Beads were added and incubated for 4 hr at 4. Beads were washed twice with each of the following buffers: wash buffer A (50mMHEPES-KOH pH 7.5, 140mMNaCl, 1mMEDTA pH 8.0, 0.1% Na-Deoxycholate, 1% Triton X-100, 0.1% SDS), wash buffer B (50 mM HEPES-KOH pH 7.9, 500 mM NaCl, 1 mM EDTA pH 8.0, 0.1% Na-Deoxycholate, 1% Triton X-100, 0.1% SDS), wash buffer C (20 mM Tris-HCl pH8.0, 250 mM LiCl, 1 mM EDTA pH 8.0, 0.5% Na-Deoxycholate, 0.5% IGEPAL C-630, 0.1% SDS), wash buffer D (TE with 0.2% Triton X-100), and TE buffer. To purify eluted DNA, 200 ul TE was added and then RNA was degraded by the addition of 2.5 µl of 33 mg/mL RNase A (Sigma, R4642) and incubation at 37°C for 2 hours. The DNA was then resuspended in 50 ul TE and amplified with the VAHTS® Universal DNA Library Prep Kit for Illumina V3 (Vazyme ND607). Amplified ChIP libraries were sequenced on the Illumina Novaseq 6000 platform.

ChIP-seq analysis was conducted as previously described^83^ with modifications. Briefly, CENH3 ChIP– seq Illumina reads (2 × 150 bp) from Chardonnay leaves were processed with fastp to remove adaptor sequences and low-quality bases (Phred+33-scaled quality < 20). Trimmed reads were aligned to the respective genome assembly using Bowtie2 with the default settings. Up to ten valid alignments were reported for each read pair and read pairs with Bowtie2-assigned MAPQ of <30 were discarded using samtools. Duplicate reads were removed by markduplicates in Picard tools with the following settings: -- REMOVE_DUPLICATES true --VALIDATION_STRINGENCY LENIENT. For retained read pairs that aligned to multiple locations, with varying alignment scores, the best alignment was selected. Alignments with two or more mismatches or having only one read in a pair were discarded. For each dataset, counts per million mapped reads (CPM) coverage values were calculated with the bamCoverage tool from deepTools (v.3.5.1)^84^. ChIP-seq broad peaks were called by MACS2 (v.2.2.7.1)^85^, then peaks of replicates were identified by IDR (v.2.0.4.2)^86^.

### Reference-based variant detection in grapevine accessions

For detection of SNPs for population genetic analysis, the NGS reads of 72 grapevine accessions (this study) and 475 accessions from Liang *et al*.^6^ were mapped to VHP-T2T hap1 reference genome with BWA mem (v0.7.17)^87^ using default parameters. Reads were sorted and duplicate reads were marked using Picard tools. The base qualities were re-calibrated with GATK 4.1.8^88^ and the variants were called using GATK HaplotypeCaller followed by joint genotyping with GenotypeGVCFs. To identify SVs from the haplotype-resolved genome assemblies, Syri (v1.6.3)^33^ was applied to the alignments of genome assemblies against VHP-T2T hap1 generated using minimap2 with parameters ‘-ax asm5 -eqx’ to detect SVs including insertions, deletions, inversions, and translocations. For pan-SV analysis, the identified SVs from each sample were merged using SURVIVOR (v1.0.7)^89^ with parameters ‘50 1 0 0 0 0’. The SVs in intergenic, upstream, downstream, intronic, and exonic regions were annotated using ANNOVAR^90^.

### Phylogenetic tree construction

High-quality SNPs were extracted from the raw variant calling output of GATK by VCFtools v0.1.16 using ‘--remove-indels --max-missing 0.3 --maf 0.05’ options^91^. Synonymous SNPs were sorted out after the SNPs were annotated by SNPeff version 5.1d^92^. In order to create the maximum likelihood (model Jukes-Cantor) tree, FastTree version 2.1.11 SSE3 was used using the default values for the large tree^93^. For the tree with 72 accessions, the SNPs were further filtered by samples and the tree was generated as described. For the haplotype-aware tree, a reference-free pangenome with 71 accessions and Chardonnay T2T was constructed using PGGB^94^ with parameters ‘ -n 144 -t 8 -p 93 -s 50000 -V ‘VHP-T2T:.:1000’ -S -m’^29^. The SNPs were filtered from the variants that were deconstructed from the pangenome using VHP-T2T hap1 as the reference. SNP tree was then built for each chromosome of all haplotype genomes and the coalescence tree was built using ASTER v1.15 and the ASTRAL-pro algorithm^30^.

### Biogeography analysis

To construct PCA plots, we used Plink v1.90b6.27^95^ with parameters ‘ --double-id --allow-extra-chr -- set-missing-var-ids @:# --indep-pairwise 50 10 0.1’ and then with parameters ‘ --double-id --allow-extra-chr --set-missing-var-ids @:# --extract vitis.prune.in --make-bed --pcà. Ancestry analysis was performed with ADMIXTURE version 1.3.0 with k=2 to k=12^96^.

### Pangenome analysis

To identify homologous relationships among the genomes of grapevines assembled in this study, the longest transcript of each predicted gene in each genome was chosen as a representative. An all-against-all comparison was then performed using BLASTP followed by clustering using OrthoFinder (v.2.5.2) with default parameters. Ortholog groups among the 144 *Vitis* genomes were identified using OrthoFinder (v2.5.5)^97^ with default parameters. Based on the clustering results, we classified genes into the following four categories: core (these were shared among all 144 haplotypes), softcore (these were present in >90% of samples but not all; 130-143), dispensable (these were present in more than one but less than 130), and private (these only existed in one accessions). The GO enrichment analysis was performed using the ClusterProfiler R package^98^. Multiple sequence alignment was performed using ParaAT (v2.0)^99^. The nucleotide diversity was calculated with nuc.div function from pegas R package. Ka/Ks ratio was calculated using KaKs_calculator 3.0.

### Transcriptome sequencing and analysis

To prepare the library for RNA sequencing, a total amount of 1-3 μg RNA per sample was used as input material for the RNA sample preparations. Sequencing libraries were generated using VAHTS Universal V6 RNA-seq Library Prep Kit for Illumina ® (NR604-01/02) following the manufacturer’s recommendations and index codes were added to attribute sequences to each sample. The library’s RNA concentration was initially quantified using the Qubit® RNA Assay Kit in Qubit® 3.0, and the concentration was subsequently diluted to 1 ng/µl. The Agilent Bioanalyzer 2100 system (Agilent Technologies, CA, USA) was used to measure the insert size. After making sure that the insert size was as anticipated, the Bio-RAD CFX 96 fluorescence quantitative PCR apparatus was used to precisely measure the library effective concentration (Library effective concentration > 10 nm). HiSeq X Ten sequencing platform (Biomarker Inc, Qingdao) was used to execute the paired-end sequencing program (PE150) to produce 150 bp sequence reads. For data analysis, fastp (v0.23.4)^100^ was used to filter the raw RNA-seq data to obtain clean reads. After that, Hisat2 (v2.2.1)^101^ was used to map clean reads to the reference genome. Based on the RNA-seq alignments, StringTie (v2.2.1)^102^ computed the FPKM, TPM, and count values for each gene’s level of expression. To find the genes that were significantly up- and down-regulated, differential gene expression analysis in R using the DESeq2 package was employed. We defined differentially expressed genes as those with an absolute fold change ≥ 2 and an adjusted p-value < 0.05. For the validation of *VvLHT8* transcription, 500 ng of RNA was used to synthesize cDNA using SPARKscript Ⅱ RT Plus Kit (With gDNA Eraser) (Sparkjade). Quantitative PCR was performed using 2×SYBR Green qPCR Mix (With ROX) (Sparkjade) and CFX Opus 96 (Bio-rad). Primers used in PCR amplification are as follows: 5′-GTGTGTCATACAGCCCACCA-3′ and 5′- TTTGAAGGCATTGTCCCCTGT-3′ for *VvLHT8*; and 5′-CTTGCATCCCTCAGCACCTT-3′ and 5′- TCCTGTGGACAATGGATGGA-3′ for *VvACT1* as a reference.

### Graph-pangenome construction and structure variant genotyping

To integrate the linear reference genome and large-scale genomic variant information, we constructed a graph-based genome of grapevine using *vg* (v.1.38.0). By combining the grapevine genomes from this work and the *V. sylvestris* genomes published from a prior study with the VHP-T2T reference genome and SVs in terms of insertions and deletions more than 50 bp, we created a graph-based genome of grape using *vg construct* (v1.43.0)^103^ without removing any alternate alleles. The preliminary graph was indexed in XG and GBWT by using *vg index* with the ‘-L’ parameter to retain alternative allele paths. A GBWT index was then built using *vg gbwt* with the ‘-P’ option. Then we mapped resequencing data (50x coverage Illumina paired-end reads) of 113 grape accessions (**Table S4**) using *vg giraffe* pipeline against the pangenome graph indexed by *vg index*. Low-quality alignments with a mapping quality and base quality both below five were excluded. Finally, SV genotyping was conducted using *vg call* with default parameters.

### eQTL analysis

The resultant vcf files of SV genotyping were imputed using Beagle v5.2^104^ with default settings. SVs with minor allele frequency ≥ 0.05 were kept by VCFtools and used for downstream analysis. PCA was performed to infer population structure using R package rMVP. Genes with a mean TPM value larger than 0.1 were contained. The top 20 hidden and confounding factors in expression data were inferred using the probabilistic estimation of expression residuals (PEER) method. Both the first 20 factors in PEER results and the first ten principal components in PCA analysis were used as covariates. The eQTL mapping was conducted using MatrixEQTL^105^. The resultant P-value were adjusted using the FDR method. The significance thresholds (*P* < 2.14×10^-5^) were calculated using the Genetic type 1 Error Calculator^106^. For each gene, eQTLs were classified as *cis* if they were located within 5 kb upstream or downstream of the gene’s annotated transcription start site or transcription stop site. Primers used for the SV validation in **Figure 6G** are as follows: 5′-CAATGGAGGTAGGCCTCGTT-3′ and 5′- TCCCCACCTTTCTCTTTGGG-3′.

### Measurement of stomatal conductance, density and size

The 4-7th fully-expended leaves from the tip of a branch were used for characterizing stomatal traits of grapevines according to our previous study^107^. A portable hand-held LI-600 fluoro-stomata measuring instrument (LI-COR, US) was used to measure the stomatal conductance on the abaxial side at 10:00 AM and 22:00 PM. For the measurement of stomatal density and size, the abaxial epidermis were imaged using a microscope (Eclipse Ci-L, Nikon, Japan), followed by measurement using Image J software (NIH, Bethesda, MD). The long axe of each stomate was measured as the size of a stomate. At least 5 leaves from different grapevines were used for all the measurement.

### Downy mildew infection assay

Downy mildew infection was performed as described previously^108^. Each grapevine leaf disc (8 mm in diameter) was inoculated with 20 μL of 1×10^5^ sporangia/mL *P. viticola* suspension on the abaxial surface and incubated at 18 ± 2°C with relative humidity of 80 ± 10% in a growth chamber under a 12 h white light (60 μmol·m^−2^·s^−1^)/12 h dark regime. Leaf discs were collected 24 hours post inoculation for transcriptome analysis and 7 days post inoculation for sporangium counting. At least three biological replicates were performed with five leaf discs from different grapevines in each replication.

### Salicylic acid quantification

Fully-expended leaves of grapevines were used for SA quantification according to the previous method^109^. All the detections were performed on a Vanquish UHPLC system combined with a TSQ Altis MS/MS system (Thermo Scientific, USA). ACQUITY UPLC@HSS T3 column (150 mm × 2.1 mm, 1.8 μm particle size, Waters) was used for the separation of samples with the column oven at 35°C. The mobile phase was composed of 0.1% formic acid in water (A) and 0.1% formic acid in ACN (B) with the flow rate at 0.2 mL/min. A linear gradient elution program with the following proportions (v/v) of solvent B was applied: 0-1 min at 10%, 1-13 min from 10% to 60%, 13-14 min from 60% to 95%, 14-17 min at 95%, 17-18.1 min from 95% to 10%, 18.1-20 min at 10%, giving a total run time of 20 min. The injection volume was 2 µL.

Salicylic acid was monitored by multiple reaction monitoring in a positive mode of electrospray ionization. The ion source conditions were as follows: positive ion capillary, 4000 V; sheath gas, 35 arb; aux gas, 10 arb; ion transfer tube temperature, 350°C; Vaporizer temperature, 350°C. After optimizing the MRM parameters, the precursor ion m/z of salicylic acid under positive mode was 136.8, as well as the main specific products ion m/z were 64.883 and 92.883 with the collision energy of 28.71 and 16.02 V, respectively. The RF lens was set at 42 V.

### Statistical analysis

Details of the statistics applied in this study are provided in the figures, figure legends and methods. All statistics were carried out in R using Student’s or Tukey’s t-test or Wilcoxon rank sum where appropriate (unless otherwise indicated).

## Supporting information

All supplementary figures

## Author contributions

L.G. and W.Y. conceived and supervised the project. J.J., W.Z., J.M., L.L.G., X.Z., H.S. and C.L. curated and prepared the grapevine samples. D.W., W.Z., J.M., W.J., J.J., G.Q., L.L.G., Q.Y., X.Z., J.W. and H.S. performed the phenotyping and molecular experiments. D.M. and Q.Y. conducted ChIP-seq experiments. D.H.A., X.W., S.C., J.S. and S.Y. conducted genome assembly and annotation. D.H.A and M.Y. performed variant calling, constructed pangenome graph and conducted eQTL mapping. L.G., D.H.A., M.S.R and W.Y. interpretated results. M.N.B. provided technical assistance and participated the discussion. L.G., D.H.A., M.S.R., X.W., M.Y. and W.Y. prepared the figures and tables, and wrote the manuscript. All authors read and approve the manuscript.

## Acknowledgement

We thank Bioinformatics Platform at Peking University Institute of Advanced Agricultural Sciences (PKU-IAAS) for providing high-performance computing resources. We thank Xiaoyu Liu and Yan Li from PKU-IAAS Mass Spectrometry Platform for their technical support in quantification of salicylic acid. Both W.Y. and L.G laboratory are supported by Taishan Scholars Program of Shandong Province and Shandong Provincial Science and Technology Innovation Fund. L.G. is also supported by Natural Science Foundation for Distinguished Young Scholars of Shandong Province (ZR2023JQ010).

## Literature cited

1. Dong, Y. et al. Dual domestications and origin of traits in grapevine evolution. Science 379, 892–901 (2023).

2. Aradhya, M. K. et al. Genetic structure and differentiation in cultivated grape, *Vitis vinifera* L. Genet. Res. 81, 179–192 (2003).

3. Terral, J.-F. et al. Evolution and history of grapevine (*Vitis vinifera*) under domestication: new morphometric perspectives to understand seed domestication syndrome and reveal origins of ancient European cultivars. Ann. Bot. 105, 443–455 (2010).

4. Grassi, F. & De Lorenzis, G. Back to the Origins: Background and Perspectives of Grapevine Domestication. Int. J. Mol. Sci. 22, 4518 (2021).

5. Walker, M. A., Heinitz, C., Riaz, S. & Uretsky, J. Grape Taxonomy and Germplasm. in The Grape Genome (eds. Cantu, D. & Walker, M. A.) 25–38 (Springer International Publishing, 2019). doi:10.1007/978-3-030-18601-2_2.

6. Liang, Z. et al. Whole-genome resequencing of 472 Vitis accessions for grapevine diversity and demographic history analyses. Nat. Commun. 10, 1190 (2019).

7. Jaillon, O. et al. The grapevine genome sequence suggests ancestral hexaploidization in major angiosperm phyla. Nature 449, 463–467 (2007).

8. Chin, C.-S. et al. Phased diploid genome assembly with single-molecule real-time sequencing. Nat. Methods 13, 1050–1054 (2016).

9. Zhou, Y. et al. The population genetics of structural variants in grapevine domestication. Nat. Plants 5, 965–979 (2019).

10. Shirasawa, K., et al. *De novo* whole-genome assembly in an interspecific hybrid table grape, ‘Shine Muscat’. DNA Res. 29, dsac040 (2022).

11. Velt, A. et al. An improved reference of the grapevine genome reasserts the origin of the PN40024 highly homozygous genotype. G3 GenesGenomesGenetics jkad067 (2023) doi:10.1093/g3journal/jkad067.

12. Shi, X. et al. The complete reference genome for grapevine ( *Vitis vinifera* L.) genetics and breeding. Hortic. Res. 10, uhad061 (2023).

13. Zhang, K. et al. The haplotype-resolved T2T genome of teinturier cultivar Yan73 reveals the genetic basis of anthocyanin biosynthesis in grapes. Hortic. Res. 10, uhad205 (2023).

14. Wang, X. et al. Telomere-to-telomere and gap-free genome assembly of a susceptible grapevine species (Thompson Seedless) to facilitate grape functional genomics. Hortic. Res. uhad260 (2023) doi:10.1093/hr/uhad260.

15. Danilevicz, M. F., Tay Fernandez, C. G., Marsh, J. I., Bayer, P. E. & Edwards, D. Plant pangenomics: approaches, applications and advancements. Curr. Opin. Plant Biol. 54, 18–25 (2020).

16. Liao, W.-W. et al. A draft human pangenome reference. Nature 617, 312–324 (2023).

17. Zhou, Y. et al. Assembly of a pangenome for global cattle reveals missing sequences and novel structural variations, providing new insights into their diversity and evolutionary history. Genome Res. 32, 1585–1601 (2022).

18. Jiang, Y.-F. et al. Pangenome obtained by long-read sequencing of 11 genomes reveal hidden functional structural variants in pigs. iScience 26, 106119 (2023).

19. Li, R. et al. A sheep pangenome reveals the spectrum of structural variations and their effects on tail phenotypes. Genome Res. 33, 463–477 (2023).

20. Liu, Y. et al. Pan-Genome of Wild and Cultivated Soybeans. Cell 182, 162–176.e13 (2020).

21. Shang, L. et al. A super pan-genomic landscape of rice. Cell Res. 32, 878–896 (2022).

22. Li, N. et al. Super-pangenome analyses highlight genomic diversity and structural variation across wild and cultivated tomato species. Nat. Genet. 55, 852–860 (2023).

23. Zhou, Y. et al. Graph pangenome captures missing heritability and empowers tomato breeding. Nature 606, 527–534 (2022).

24. Cochetel, N. et al. A super-pangenome of the North American wild grape species. Genome Biol. 24, 290 (2023).

25. Cheng, H. et al. Haplotype-resolved assembly of diploid genomes without parental data. Nat. Biotechnol. 40, 1332–1335 (2022).

26. Dudchenko, O. et al. De novo assembly of the Aedes aegypti genome using Hi-C yields chromosome-length scaffolds. Science 356, 92–95 (2017).

27. Shi, X. et al. The complete reference genome for grapevine (*Vitis vinifera* L.) genetics and breeding. Hortic. Res. 10, uhad061 (2023).

28. Cochetel, N. et al. Diploid chromosome-scale assembly of the *Muscadinia rotundifolia* genome supports chromosome fusion and disease resistance gene expansion during *Vitis* and *Muscadinia* divergence. G3 GenesGenomesGenetics 11, jkab033 (2021).

29. Garrison, E. et al. Building pangenome graphs. 2023.04.05.535718 Preprint at 10.1101/2023.04.05.535718 (2023).

30. Zhang, C., Scornavacca, C., Molloy, E. K. & Mirarab, S. ASTRAL-Pro: Quartet-Based Species-Tree Inference despite Paralogy. Mol. Biol. Evol. 37, 3292–3307 (2020).

31. Marone, D., Russo, M., Laidò, G., De Leonardis, A. & Mastrangelo, A. Plant Nucleotide Binding Site– Leucine-Rich Repeat (NBS-LRR) Genes: Active Guardians in Host Defense Responses. Int. J. Mol. Sci. 14, 7302–7326 (2013).

32. Steuernagel, B. et al. The NLR-Annotator Tool Enables Annotation of the Intracellular Immune Receptor Repertoire. Plant Physiol. 183, 468–482 (2020).

33. Goel, M., Sun, H., Jiao, W.-B. & Schneeberger, K. SyRI: finding genomic rearrangements and local sequence differences from whole-genome assemblies. Genome Biol. 20, 277 (2019).

34. Foria, S. et al. InDel markers for monitoring the introgression of downy mildew resistance from wild relatives into grape varieties. Mol. Breed. 38, 124 (2018).

35. Di Gaspero, G. et al. Selective sweep at the Rpv3 locus during grapevine breeding for downy mildew resistance. Theor. Appl. Genet. 124, 277–286 (2012).

36. Zhou, H. et al. Patellin protein family functions in plant development and stress response. J. Plant Physiol. 234–235, 94–97 (2019).

37. He, Q. et al. A graph-based genome and pan-genome variation of the model plant Setaria. Nat. Genet. 55, 1232–1242 (2023).

38. Sirén, J. et al. Pangenomics enables genotyping of known structural variants in 5202 diverse genomes. Science 374, abg8871 (2021).

39. GTEx Consortium et al. The impact of structural variation on human gene expression. Nat. Genet. 49, 692–699 (2017).

40. Liu, G. et al. Amino Acid Homeostasis Modulates Salicylic Acid–Associated Redox Status and Defense Responses in *Arabidopsis*. Plant Cell 22, 3845–3863 (2010).

41. Murata, Y., Mori, I. C. & Munemasa, S. Diverse Stomatal Signaling and the Signal Integration Mechanism. Annu. Rev. Plant Biol. 66, 369–392 (2015).

42. Coupel-Ledru, A. et al. Reduced nighttime transpiration is a relevant breeding target for high water-use efficiency in grapevine. Proc. Natl. Acad. Sci. 113, 8963–8968 (2016).

43. Ye, W. et al. Stomatal immunity against fungal invasion comprises not only chitin-induced stomatal closure but also chitosan-induced guard cell death. Proc. Natl. Acad. Sci. 117, 20932–20942 (2020).

44. Alston, J. M. & Sambucci, O. Grapes in the World Economy. in The Grape Genome (eds. Cantu, D. & Walker, M. A.) 1–24 (Springer International Publishing, 2019). doi:10.1007/978-3-030-18601-2_1.

45. Minio, A., Cochetel, N., Massonnet, M., Figueroa-Balderas, R. & Cantu, D. HiFi chromosome-scale diploid assemblies of the grape rootstocks 110R, Kober 5BB, and 101–14 Mgt. Sci. Data 9, 660 (2022).

46. Zou, C., et al. Multiple independent recombinations led to hermaphroditism in grapevine. Proc. Natl. Acad. Sci. 118, e2023548118 (2021).

47. Massonnet, M. et al. The genetic basis of sex determination in grapes. Nat. Commun. 11, 2902 (2020).

48. Chin, C.-S. et al. Phased diploid genome assembly with single-molecule real-time sequencing. Nat. Methods 13, 1050–1054 (2016).

49. Blanco-Ulate, B., Allen, G., Powell, A. L. T. & Cantu, D. Draft Genome Sequence of *Botrytis cinerea* BcDW1, Inoculum for Noble Rot of Grape Berries. Genome Announc. 1, e00252–13 (2013).

50. Velasco, R. et al. A High Quality Draft Consensus Sequence of the Genome of a Heterozygous Grapevine Variety. PLoS ONE 2, e1326 (2007).

51. Porebski, S., Bailey, L. G. & Baum, B. R. Modification of a CTAB DNA extraction protocol for plants containing high polysaccharide and polyphenol components. Plant Mol. Biol. Report. 15, 8–15 (1997).

52. Belton, J.-M. et al. Hi–C: A comprehensive technique to capture the conformation of genomes. Methods 58, 268–276 (2012).

53. Servant, N. et al. HiC-Pro: an optimized and flexible pipeline for Hi-C data processing. Genome Biol. 16, 259 (2015).

54. Marçais, G. & Kingsford, C. A fast, lock-free approach for efficient parallel counting of occurrences of k-mers. Bioinforma. Oxf. Engl. 27, 764–770 (2011).

55. Vurture, G. W. et al. GenomeScope: fast reference-free genome profiling from short reads. Bioinformatics 33, 2202–2204 (2017).

56. Cheng, H., Concepcion, G. T., Feng, X., Zhang, H. & Li, H. Haplotype-resolved de novo assembly using phased assembly graphs with hifiasm. Nat. Methods 18, 170–175 (2021).

57. Durand, N. C. et al. Juicer Provides a One-Click System for Analyzing Loop-Resolution Hi-C Experiments. Cell Syst. 3, 95–98 (2016).

58. Dudchenko, O. et al. De novo assembly of the *Aedes aegypti* genome using Hi-C yields chromosome-length scaffolds. Science 356, 92–95 (2017).

59. Langmead, B. & Salzberg, S. L. Fast gapped-read alignment with Bowtie 2. Nat. Methods 9, 357–359 (2012).

60. Xu, M. et al. TGS-GapCloser: A fast and accurate gap closer for large genomes with low coverage of error-prone long reads. GigaScience 9, giaa094 (2020).

61. Durand, N. C. et al. Juicebox Provides a Visualization System for Hi-C Contact Maps with Unlimited Zoom. Cell Syst. 3, 99–101 (2016).

62. Li, H. Minimap2: pairwise alignment for nucleotide sequences. Bioinformatics 34, 3094–3100 (2018).

63. Jain, C., Rhie, A., Hansen, N. F., Koren, S. & Phillippy, A. M. Long-read mapping to repetitive reference sequences using Winnowmap2. Nat. Methods 19, 705–710 (2022).

64. Poplin, R. et al. A universal SNP and small-indel variant caller using deep neural networks. Nat. Biotechnol. 36, 983–987 (2018).

65. Li, H. A statistical framework for SNP calling, mutation discovery, association mapping and population genetical parameter estimation from sequencing data. Bioinformatics 27, 2987–2993 (2011).

66. Li, H. et al. The Sequence Alignment/Map format and SAMtools. Bioinformatics 25, 2078–2079 (2009).

67. Manni, M., Berkeley, M. R., Seppey, M. & Zdobnov, E. M. BUSCO: Assessing Genomic Data Quality and Beyond. Curr. Protoc. 1, e323 (2021).

68. Ou, S. & Jiang, N. LTR_retriever: A Highly Accurate and Sensitive Program for Identification of Long Terminal Repeat Retrotransposons. Plant Physiol. 176, 1410–1422 (2018).

69. Rhie, A., Walenz, B. P., Koren, S. & Phillippy, A. M. Merqury: reference-free quality, completeness, and phasing assessment for genome assemblies. Genome Biol. 21, 245 (2020).

70. Flynn, J. M. et al. RepeatModeler2 for automated genomic discovery of transposable element families. Proc. Natl. Acad. Sci. U. S. A. 117, 9451–9457 (2020).

71. Nishimura, D. RepeatMasker. Biotech Softw. Internet Rep. 1, 36–39 (2000).

72. Brůna, T., Hoff, K. J., Lomsadze, A., Stanke, M. & Borodovsky, M. BRAKER2: automatic eukaryotic genome annotation with GeneMark-EP+ and AUGUSTUS supported by a protein database. NAR Genomics Bioinforma. 3, lqaa108 (2021).

73. Johnson, A. D. et al. SNAP: a web-based tool for identification and annotation of proxy SNPs using HapMap. Bioinforma. Oxf. Engl. 24, 2938–2939 (2008).

74. Cantarel, B. L. et al. MAKER: an easy-to-use annotation pipeline designed for emerging model organism genomes. Genome Res. 18, 188–196 (2008).

75. Massonnet, M. et al. The genetic basis of sex determination in grapes. Nat. Commun. 11, 2902 (2020).

76. Minio, A., Cochetel, N., Massonnet, M., Figueroa-Balderas, R. & Cantu, D. HiFi chromosome-scale diploid assemblies of the grape rootstocks 110R, Kober 5BB, and 101–14 Mgt. Sci. Data 9, 660 (2022).

77. Fu, L., Niu, B., Zhu, Z., Wu, S. & Li, W. CD-HIT: accelerated for clustering the next-generation sequencing data. Bioinformatics 28, 3150–3152 (2012).

78. Grabherr, M. G. et al. Full-length transcriptome assembly from RNA-Seq data without a reference genome. Nat. Biotechnol. 29, 644–652 (2011).

79. Benson, G. Tandem repeats finder: a program to analyze DNA sequences. Nucleic Acids Res. 27, 573– 580 (1999).

80. Wlodzimierz, P., Hong, M. & Henderson, I. R. TRASH: Tandem Repeat Annotation and Structural Hierarchy. Bioinformatics 39, btad308 (2023).

81. Huang, H.-R. et al. Telomere-to-telomere haplotype-resolved reference genome reveals subgenome divergence and disease resistance in triploid Cavendish banana. Hortic. Res. 10, uhad153 (2023).

82. Vollger, M. R., Kerpedjiev, P., Phillippy, A. M. & Eichler, E. E. StainedGlass: interactive visualization of massive tandem repeat structures with identity heatmaps. Bioinformatics 38, 2049–2051 (2022).

83. Wlodzimierz, P. et al. Cycles of satellite and transposon evolution in Arabidopsis centromeres. Nature 618, 557–565 (2023).

84. Ramírez, F., Dündar, F., Diehl, S., Grüning, B. A. & Manke, T. deepTools: a flexible platform for exploring deep-sequencing data. Nucleic Acids Res. 42, W187–W191 (2014).

85. Zhang, Y. et al. Model-based Analysis of ChIP-Seq (MACS). Genome Biol. 9, R137 (2008).

86. Li, Q., Brown, J. B., Huang, H. & Bickel, P. J. Measuring reproducibility of high-throughput experiments. Ann. Appl. Stat. 5, (2011).

87. Li, H. & Durbin, R. Fast and accurate short read alignment with Burrows–Wheeler transform. Bioinformatics 25, 1754–1760 (2009).

88. Van der Auwera, G. A. et al. From FastQ data to high confidence variant calls: the Genome Analysis Toolkit best practices pipeline. Curr. Protoc. Bioinforma. 43, 11.10.1–11.10.33 (2013).

89. Jeffares, D. C. et al. Transient structural variations have strong effects on quantitative traits and reproductive isolation in fission yeast. Nat. Commun. 8, 14061 (2017).

90. Wang, K., Li, M. & Hakonarson, H. ANNOVAR: functional annotation of genetic variants from high-throughput sequencing data. Nucleic Acids Res. 38, e164–e164 (2010).

91. Danecek, P. et al. The variant call format and VCFtools. Bioinforma. Oxf. Engl. 27, 2156–2158 (2011).

92. Cingolani, P. et al. A program for annotating and predicting the effects of single nucleotide polymorphisms, SnpEff: SNPs in the genome of Drosophila melanogaster strain w 1118 ; iso-2; iso-3. Fly (Austin) 6, 80–92 (2012).

93. Price, M. N., Dehal, P. S. & Arkin, A. P. FastTree 2 – Approximately Maximum-Likelihood Trees for Large Alignments. PLoS ONE 5, e9490 (2010).

94. Garrison, E. et al. Building pangenome graphs. (2023) doi:10.1101/2023.04.05.535718.

95. Chang, C. C. et al. Second-generation PLINK: rising to the challenge of larger and richer datasets. GigaScience 4, 7 (2015).

96. Alexander, D. H., Novembre, J. & Lange, K. Fast model-based estimation of ancestry in unrelated individuals. Genome Res. 19, 1655–1664 (2009).

97. Emms, D. M. & Kelly, S. OrthoFinder: phylogenetic orthology inference for comparative genomics. Genome Biol. 20, 238 (2019).

98. Wu, T. et al. clusterProfiler 4.0: A universal enrichment tool for interpreting omics data. The Innovation 2, 100141 (2021).

99. Zhang, Z. et al. ParaAT: A parallel tool for constructing multiple protein-coding DNA alignments. Biochem. Biophys. Res. Commun. 419, 779–781 (2012).

100. Chen, S., Zhou, Y., Chen, Y. & Gu, J. fastp: an ultra-fast all-in-one FASTQ preprocessor. Bioinformatics 34, i884–i890 (2018).

101. Kim, D., Paggi, J. M., Park, C., Bennett, C. & Salzberg, S. L. Graph-based genome alignment and genotyping with HISAT2 and HISAT-genotype. Nat. Biotechnol. 37, 907–915 (2019).

102. Pertea, M. et al. StringTie enables improved reconstruction of a transcriptome from RNA-seq reads. Nat. Biotechnol. 33, 290–295 (2015).

103. Hickey, G. et al. Genotyping structural variants in pangenome graphs using the vg toolkit. Genome Biol. 21, 35 (2020).

104. Browning, B. L., Zhou, Y. & Browning, S. R. A One-Penny Imputed Genome from Next-Generation Reference Panels. Am. J. Hum. Genet. 103, 338–348 (2018).

105. Shabalin, A. A. Matrix eQTL: ultra fast eQTL analysis via large matrix operations. Bioinformatics 28, 1353–1358 (2012).

106. Li, M.-X., Yeung, J. M. Y., Cherny, S. S. & Sham, P. C. Evaluating the effective numbers of independent tests and significant p-value thresholds in commercial genotyping arrays and public imputation reference datasets. Hum. Genet. 131, 747–756 (2012).

107. Zhang, M. et al. Plasma membrane H+-ATPase overexpression increases rice yield via simultaneous enhancement of nutrient uptake and photosynthesis. Nat. Commun. 12, 735 (2021).

108. Ma, T., et al. *Plasmopara viticola* effector PvRXLR111 stabilizes VvWRKY40 to promote virulence. Mol. Plant Pathol. 22, 231–242 (2021).

109. Yao, X., Xia, N., Meng, X., Duan, C. & Pan, Q. A One-Step Polyphenol Removal Approach for Detection of Multiple Phytohormones from Grape Berry. Horticulturae 8, 548 (2022).

