## Supplementary material for "Super Pangenome of Grapevines Empowers Improvement of the Oldest Domesticated Fruit": All supplementary figures

**Figure S1. Genome assembly and validation of haplotype-resolved complete genome of *Vitis vinifera* cv. Chardonnay**

A. Genome survey of winegrape cultivar Chardonnay based on kmer frequency analysis.

B. Chromatin interaction heatmap of two T2T haploid genomes using Hi-C data.

C-D. The plot of LAI (LTR assembly index) for all chromosomes in two haploid genomes.

E. Whole-genome sequence alignment of two T2T haploid genomes (horizontal) against a published genome PN40024 (vertical).

**Figure S2. Centromere sequence alignment between haplotypes and among chromosomes**

A. Sequence identity heatmap of centromere sequence alignment across 19 chromosomes, showing the conservation and divergence of centromeres among chromosomes.

B. Sequence identity heatmap of centromere sequence alignment between two haplotypes. Representative examples of three chromosomes are provided.

**Figure S3. Population genomic analysis of grapevine accessions**

A. Maximum likelihood phylogenetic tree of 591 accessions constructed using 341,450 synonymous mutations (475 accessions from Liang et al, 2019; 116 accessions highlighted in bold, this study).

B. Box plot showing differences in stomatal density, length, and conductance in 71 accessions.

C. Cross-validation error plot for k=2 to 8 for 72 samples ADMIXTURE ancestry plot in Figure 2F.

D. ADMIXTURE ancestry plot for 591 samples based on SNPs.

E. Principal component analysis plot of 591 samples based on SNPs. 71 accessions are shown in purple (Muscadine), blue (North American), green (East Asian), and red (European).

**Figure S4. Haplotype-resolved genome assembly of 71 grapevine accessions**

A. Scatter plots showing two haplotypes of all 71 grapevine accessions for BUSCO, LAI, and QV values.

B. Box plot of chromosome size distributions of 71 grapevine accessions.

C. Total length of transposable element contents (Mb) of different classes in 71 grapevine accessions. LTR: long terminal repeats. LINE: long interspersed nuclear elements, SINE: short interspersed nuclear elements, DNA: DNA transposons.

**Figure S5. Pangenome analysis, functional enrichment and Pan-NLR (Nucleotide-binding site leucine-rich repeat) analysis**

A. Comparison of different groups of grapevine accessions for the content of softcore, dispensable and private genomes.

B-C. Nucleotide diversity (B), dN/dS (C) of core, softcore, dispensable genes.

D-G. GO enrichment of pangenome core genes (D) Softcore genes (E), dispensable genes (F) and private genes (G) for 144 haplotype grapevine genomes.

H. Variation of NLR gene families in the pan- and core-NLR with additional grapevine genomes.

I. Compositions of NLR families in the grapevine pangenome and individual genome. The histogram shows the number of NLR gene families in the 144 haploid genomes with different frequencies where colors of histograms correspond to core, softcore, dispensable, and private NLR genes. The pie chart shows the proportion of the NLR gene family marked by each composition.

Different letters in A-C indicate statistical significance ($P < 0.05$, Kruskal–Wallis test followed by the Nemenyi test).

**Figure S6. Whole-genome sequence alignment of 144 haplotype grapevine genomes (including T2T genome as reference) shown in Syri plots showing different types of structural variants and syntenic regions**

For ease of observation, only intra-chromosomal structural variants were displayed.

**Figure S7. Structural variants in grapevines and Pan-SV analysis**

A. Distribution of inversion events in terms of their distance to centromeres.

B. Distribution of translocation events in terms of their distance to centromeres.

C-F. Examples of structural variants in grapevine accessions were identified both in whole-genome alignment (left) and Hi-C interaction heatmap (Right). For each variant, the top is a reference vs. reference alignment and the bottom is a reference vs. sample alignment. Structural variants were marked using rectangles or dotted circles.

**Figure S8. Expression and phenotype of UDV305 marker and nearby SVs**
Stomatal density, stomatal length, stomatal conductance at 10 AM and 10 PM of accessions with or without SVs on (A) and near (B-C) UDV305 marker. Purple: muscadine, Blue: North American, Green: East Asian, Red: European accessions. *$P < 0.05$, Wilcoxon Rank Sum test.

**Figure S9. Expression and phenotype of UDV737 and INDEL 26032 markers and nearby SVs**
Gene expression during mock and *P. viticola* (*Pv*) infection, sporangium number at 7 dpi, stomatal density, stomatal length, stomatal conductance at 10 AM and 10 PM of accessions with or without SVs near the marker UDV737 (A) and on the marker INDEL26032 (B). Purple: muscadine, Blue: North American, Green: East Asian, Red: European accessions. *$P < 0.05$, Wilcoxon Rank Sum test.

**Figure S10. Expression and phenotype of top genes identified in eQTL analysis**
Gene expression during mock and *P. viticola* (*Pv*) infection, sporangium number at 1 dpi, stomatal density, stomatal length, stomatal conductance at 10 AM and 10 PM of accessions with or without top SVs detected by eQTL. Purple: muscadine, Blue: North American, Green: East Asian, Red: European accessions. *$P < 0.05$, Wilcoxon Rank Sum test.

**Supplementary Tables**

**Table S1**. Summary statistics of telomeres assembled in *Vitis vinifera* cv. Chardonnay VHP-T2T genome.

**Table S2**. Characterization of centromere on each chromosome identified by CENH3 ChIP-seq.

**Table S3**. Information of 71 grapevine accessions used for de novo genome assembly.

**Table S4.** Stomatal traits and downy mildew infection phenotype of *Vitis* accessions used for transcriptome analysis.

**Table S5**. Summary statistics of genome sequencing reads (PacBio HiFi, NGS, ONT and Hi-C) generated for the 72 *Vitis* accessions.

**Table S6.** Predicted genome sizes for 71 accessions based on kmer frequency analysis of short reads.

**Table S7.** Summary statistics for chromosome-level haplotype-resolved genome assembly, annotation, quality assessment for 71 grapevine accessions (142 haplotypes).

**Table S8.** The comparison of two phylogenetic trees for 71 grapevine accessions, constructed either using reference-based SNPs (diploid-based) or graph-based coalescent approach (haplotype-resolved).

**Table S9.** Summary of super pangenome gene families identified in 144 grapevine genomes. The numbers represent the count of gene members for each gene family.

**Table S10.** Summary of NLR gene families identified in 144 grapevine genomes. The numbers represent the count of NLR gene members for each gene family.

**Table S11.** Summary of expression QTLs significantly associated with downy mildew resistance of *Vitis* accessions.

**Table S12.** Summary of transcriptome datasets used for genome annotation in this study.

**Figure S1. Genome assembly and validation of haplotype-resolved complete genome of *Vitis vinifera* cv. Chardonnay**

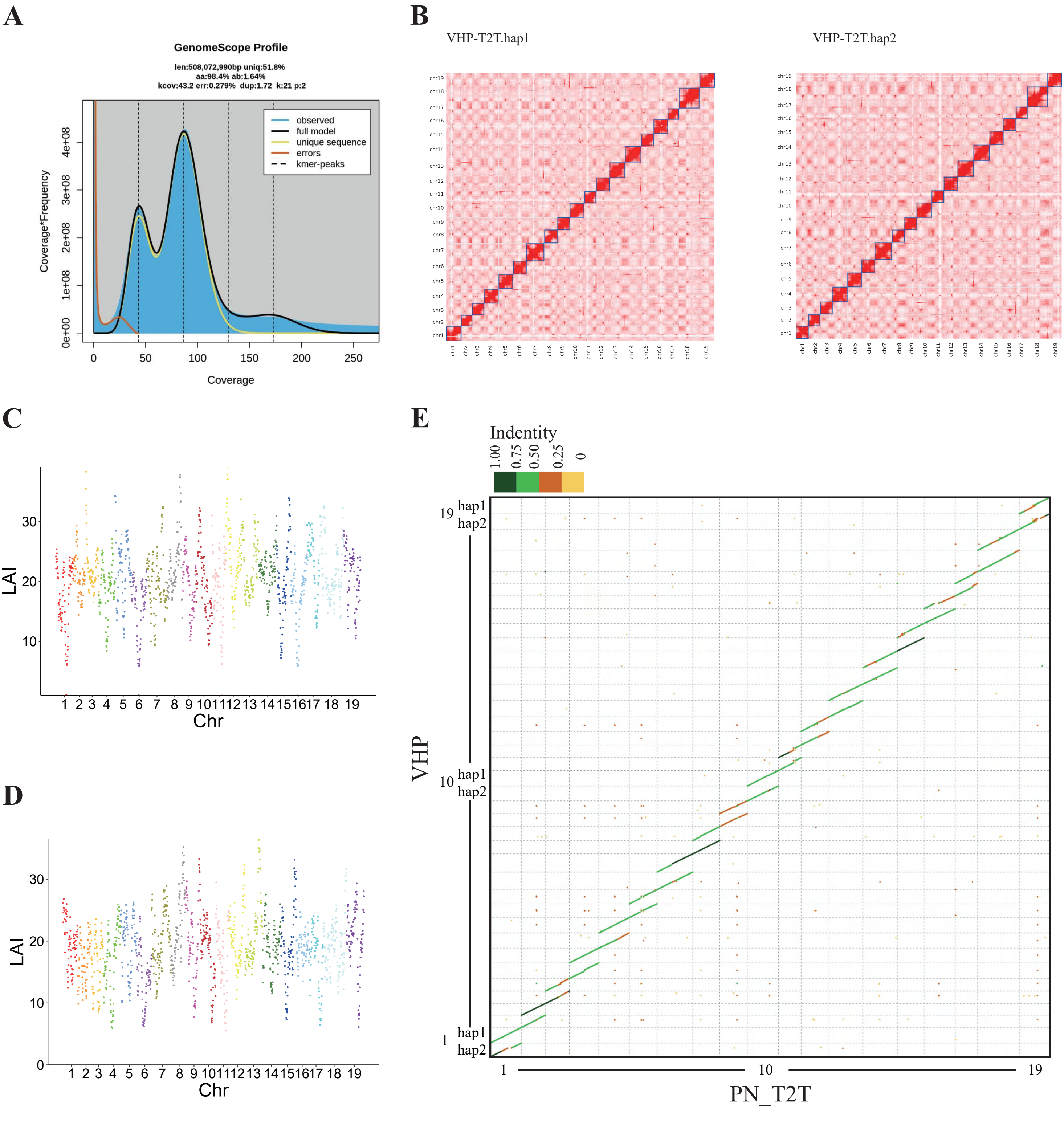

### Figure S2. Centromere sequence alignment between haplotypes and among chromosomes

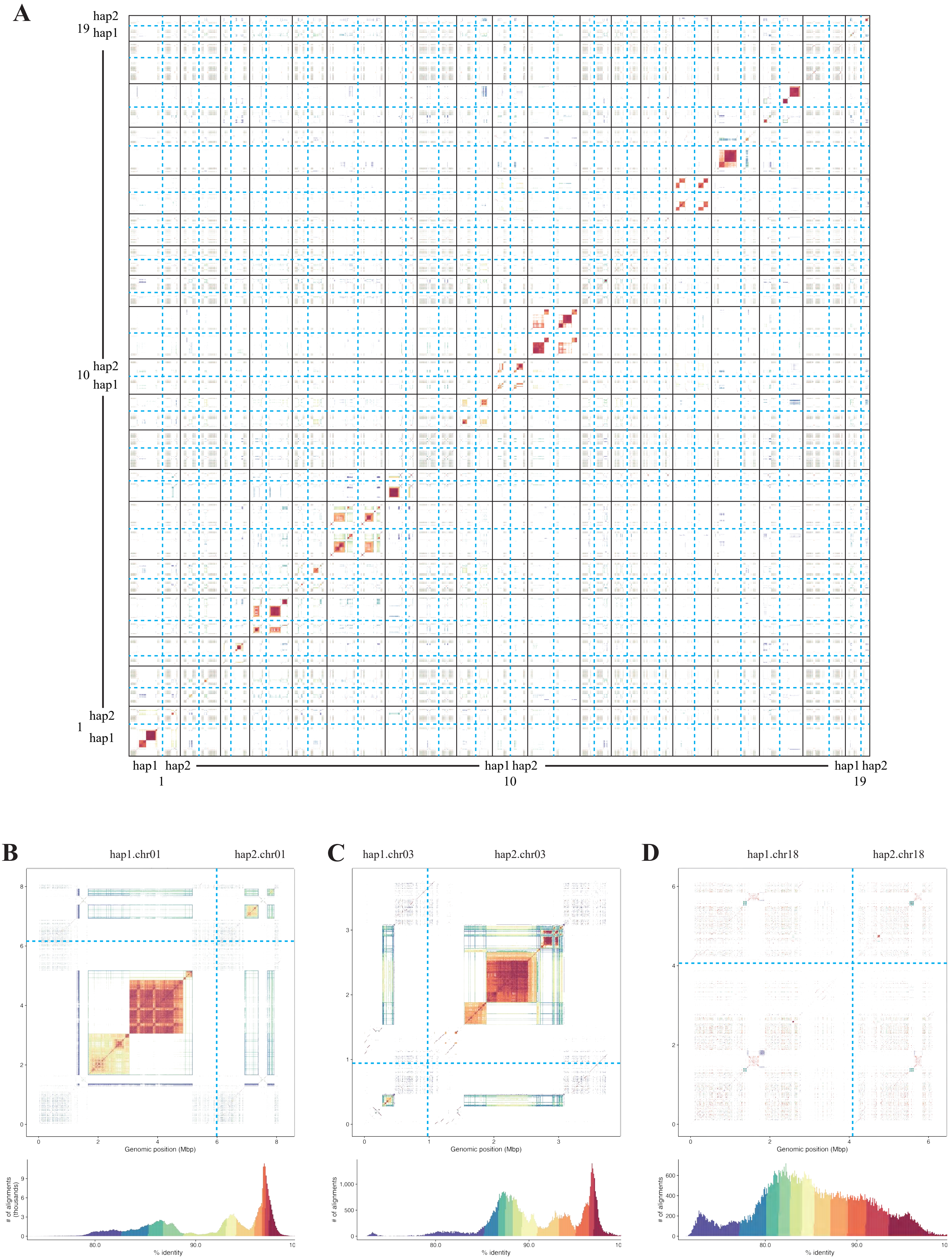

### Figure S3. Population genomic analysis of grapevine accessions

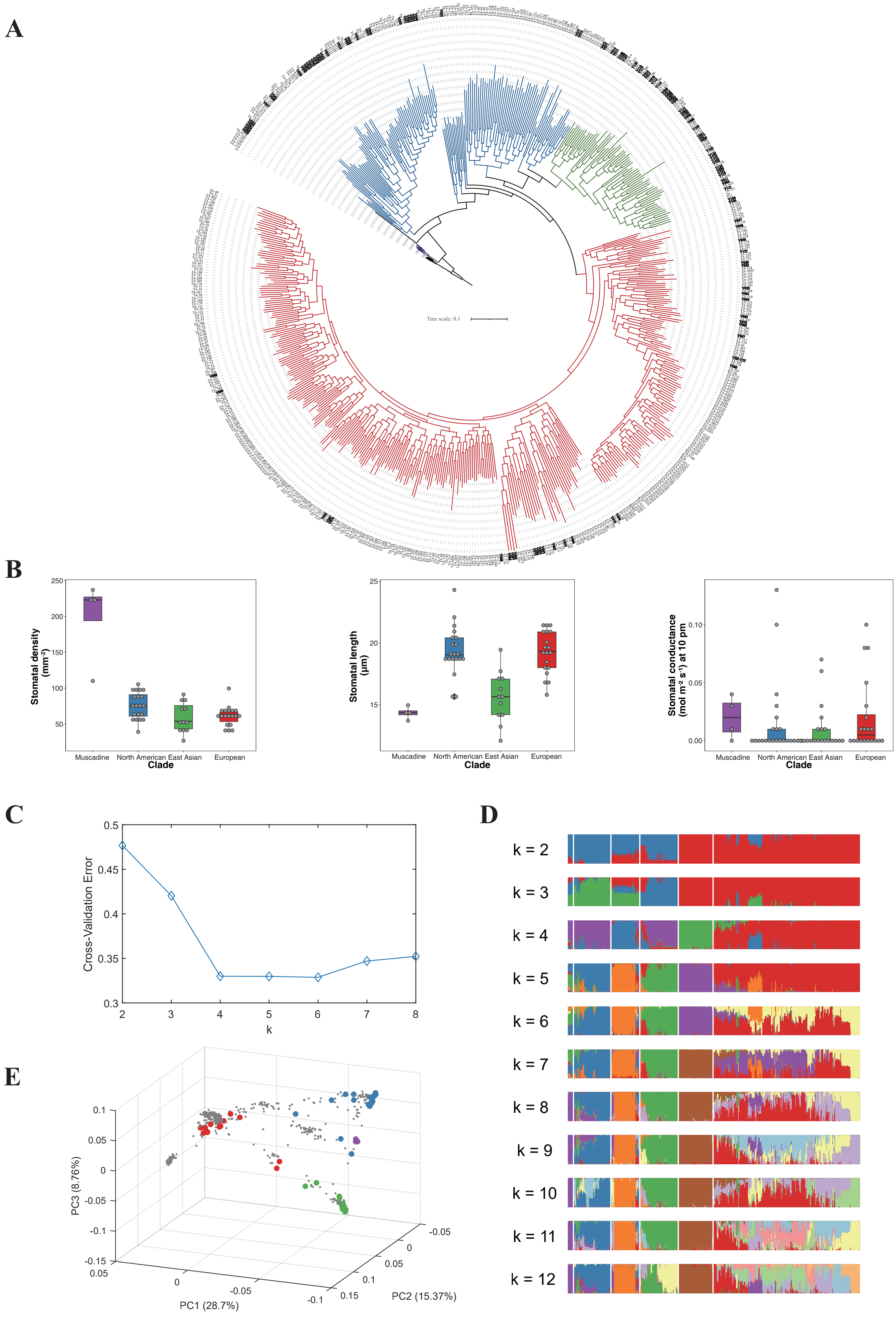

#### Figure S4. Haplotype-resolved genome assembly of 71 grapevine accessions

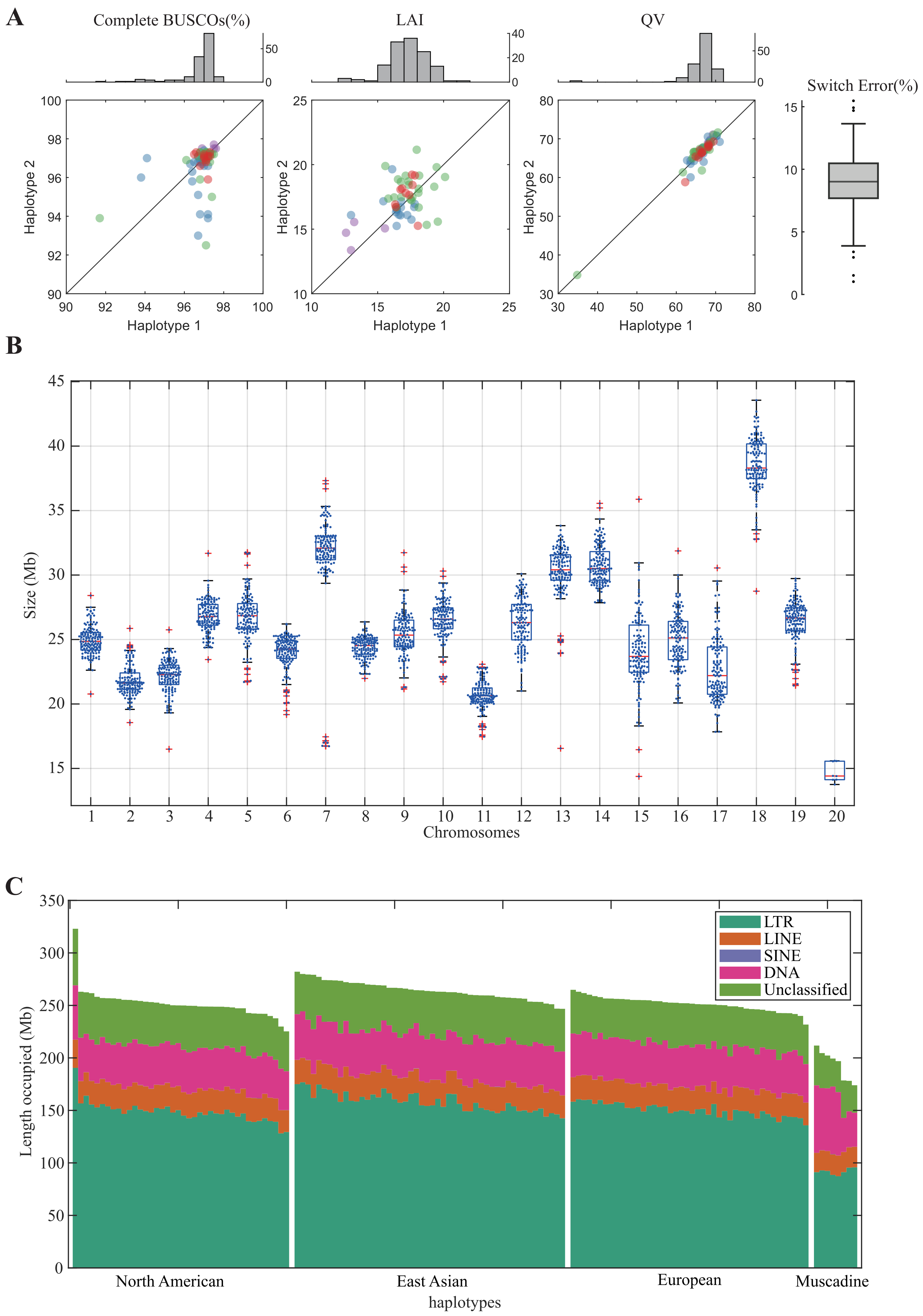

### Figure S5. Pangenome analysis, functional enrichment and Pan-NLR (Nucleotide-binding site leucine-rich repeat) analysis

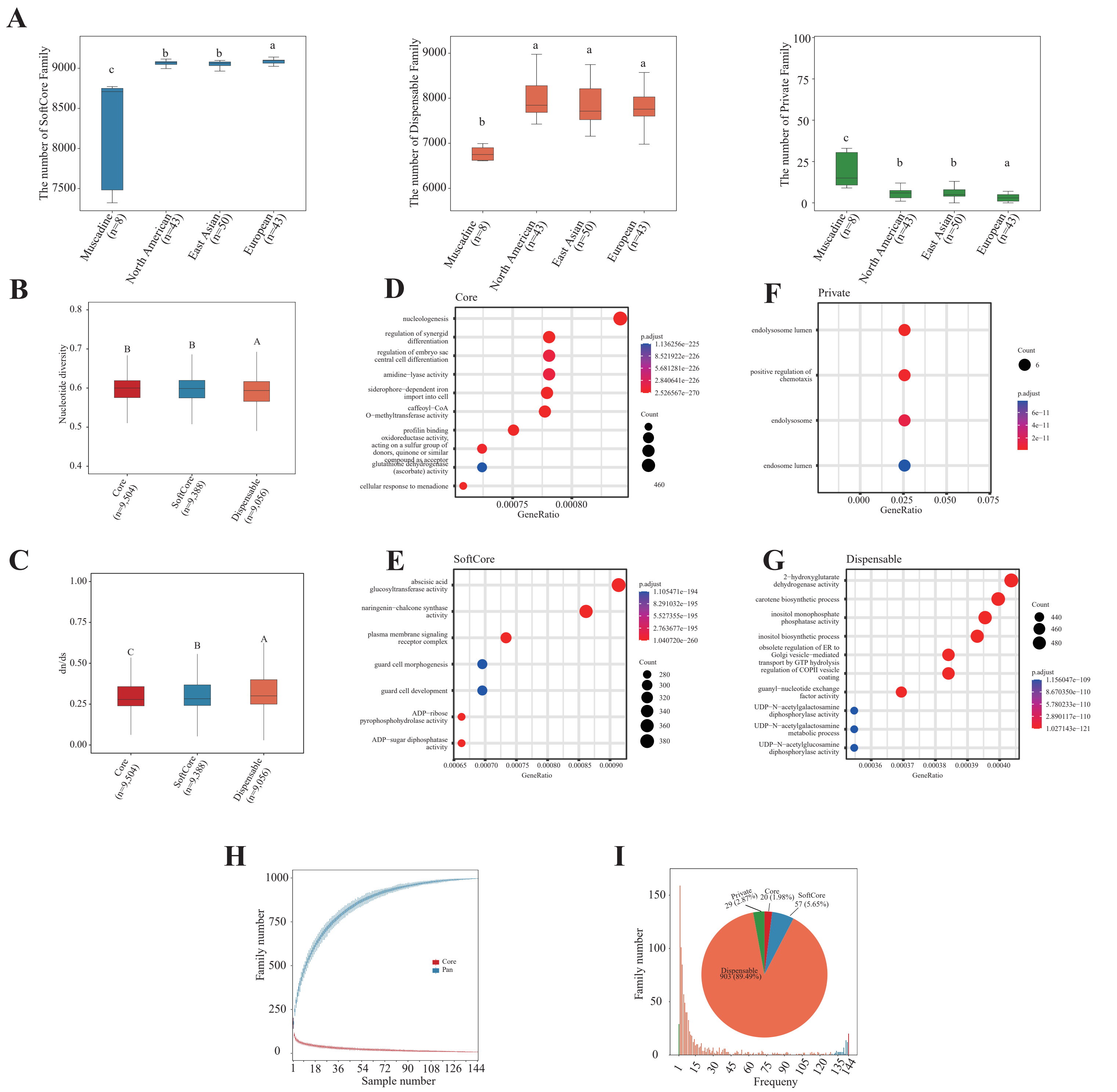

**Figure S6. Whole-genome sequence alignment of 144 haplotype grapevine genomes (including T2T genome as reference) shown in Syri plots showing different types of structural variants and syntenic regions.**

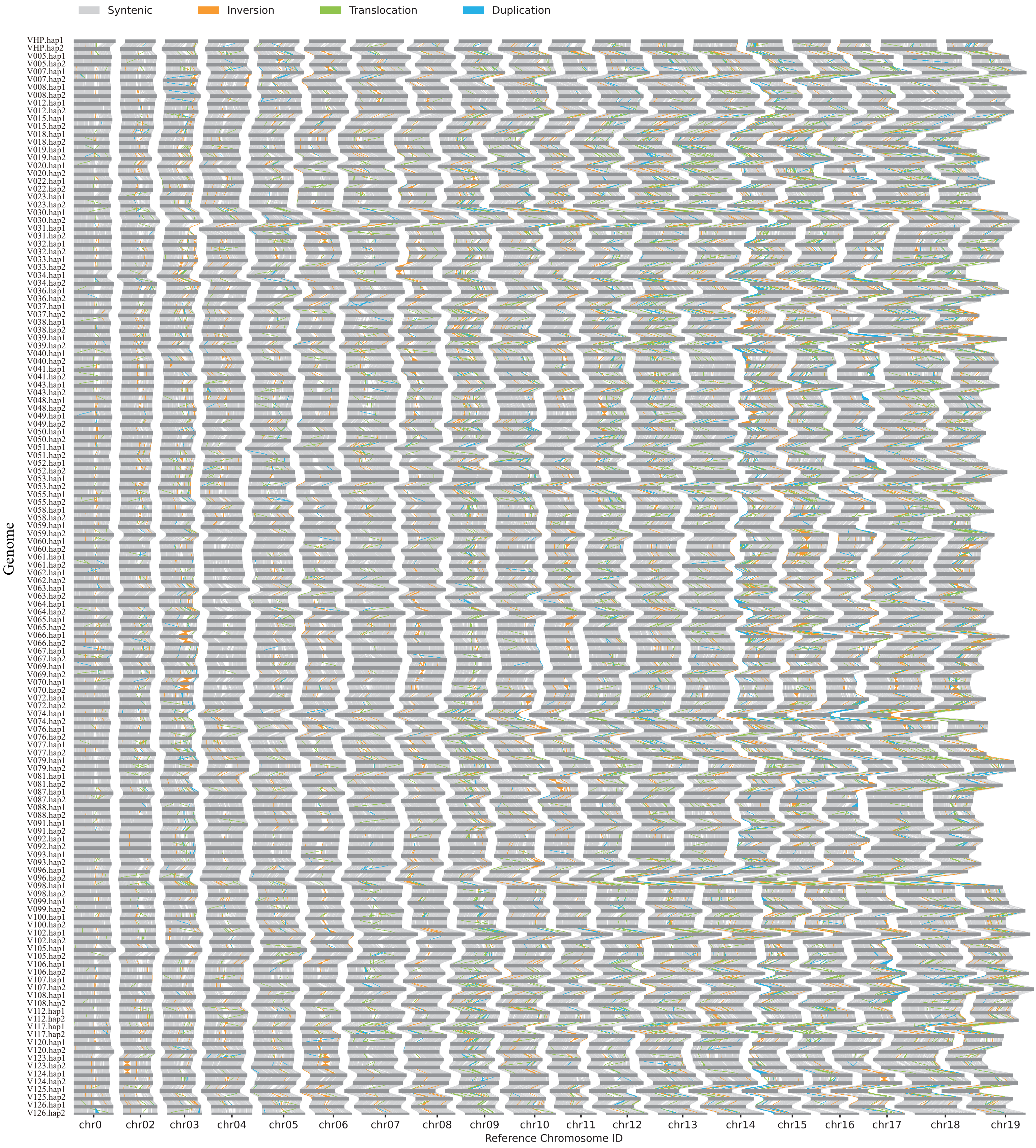

### Figure S7. Structural variants in grapevines and Pan-SV analysis

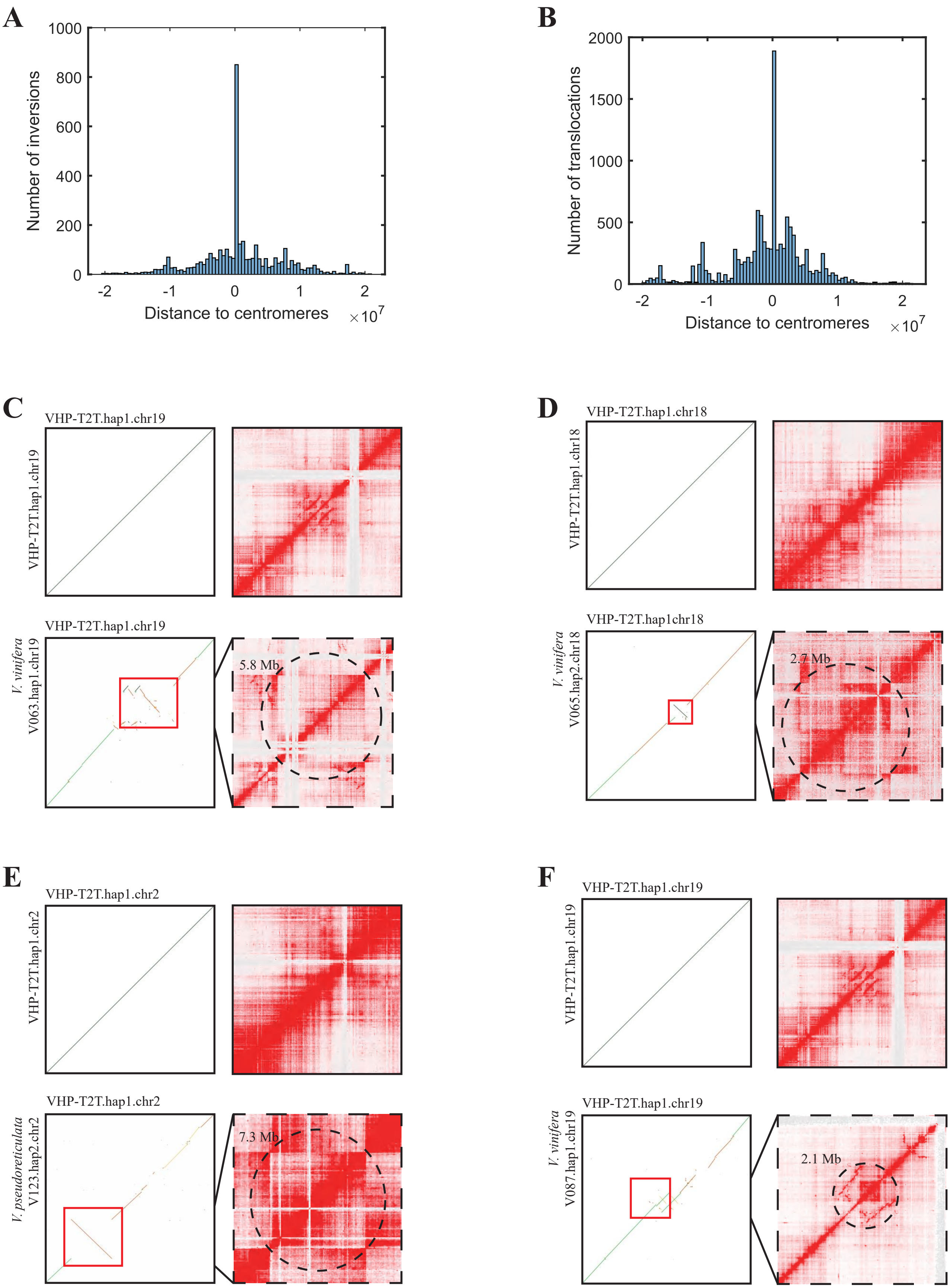

**Figure S8. Expression and phenotype of UDV305 marker and nearby SVs**

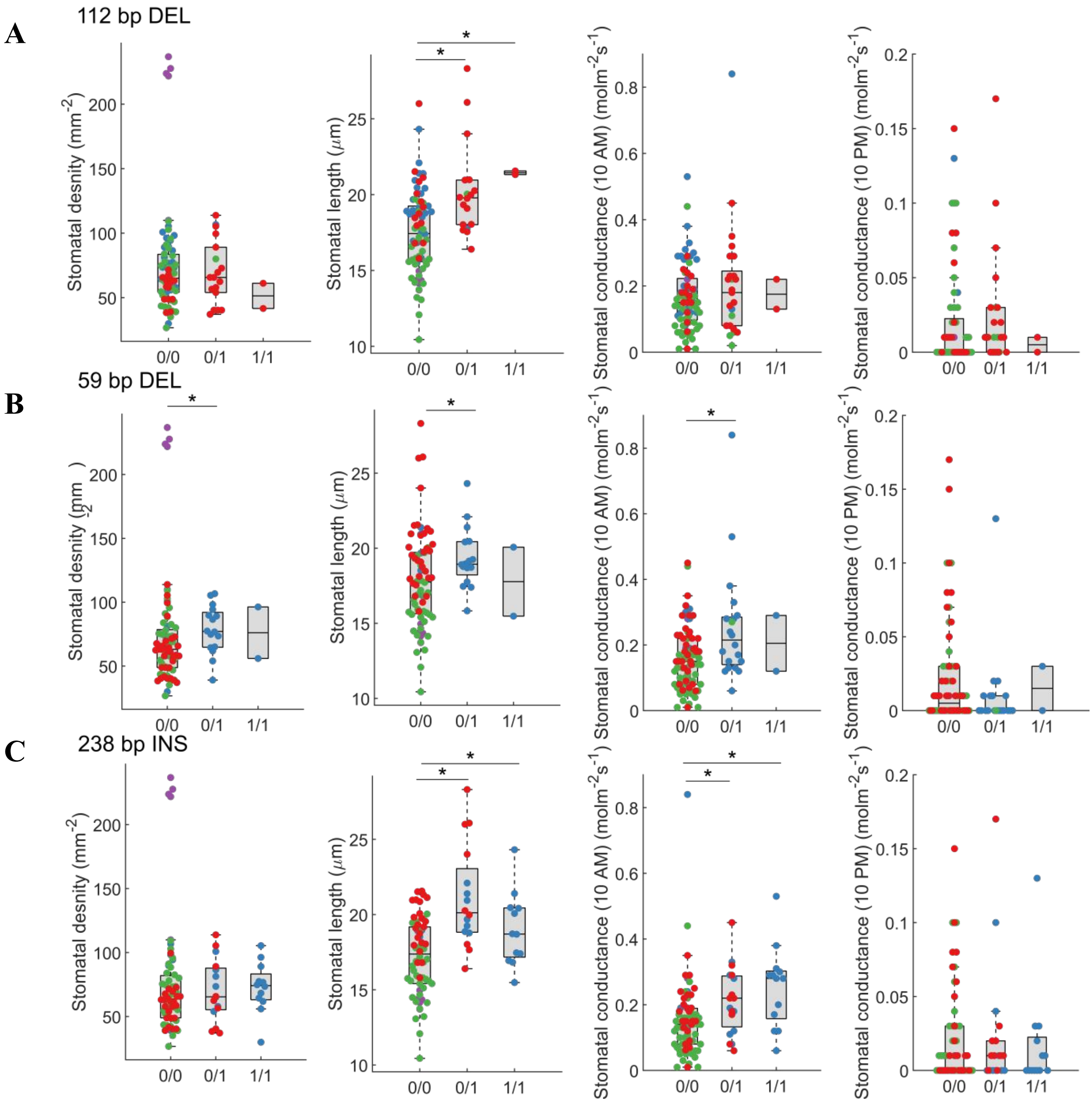

**Figure S9. Expression and phenotype of UDV737 and INDEL 26032 markers and nearby SVs**

**A**

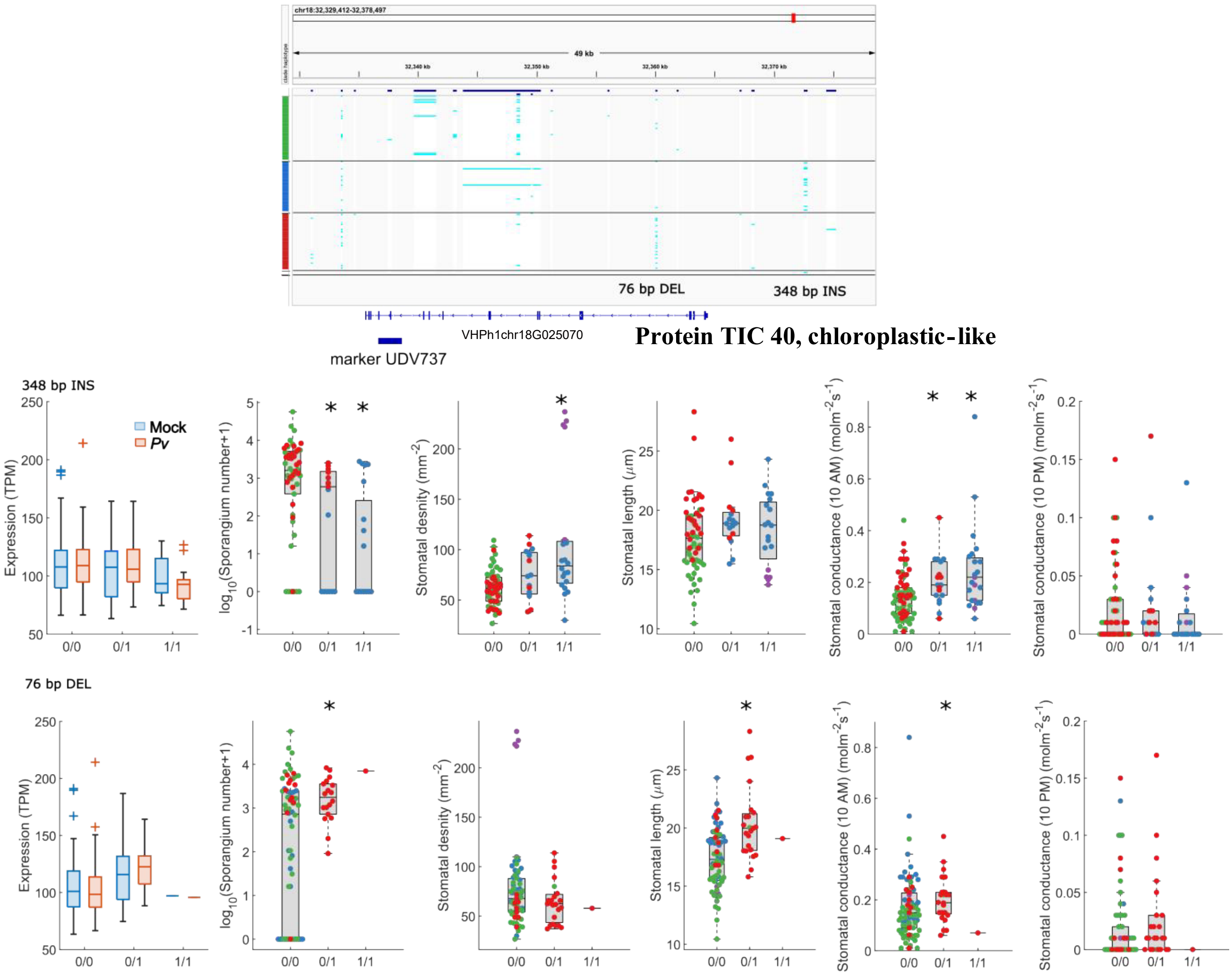


**B**

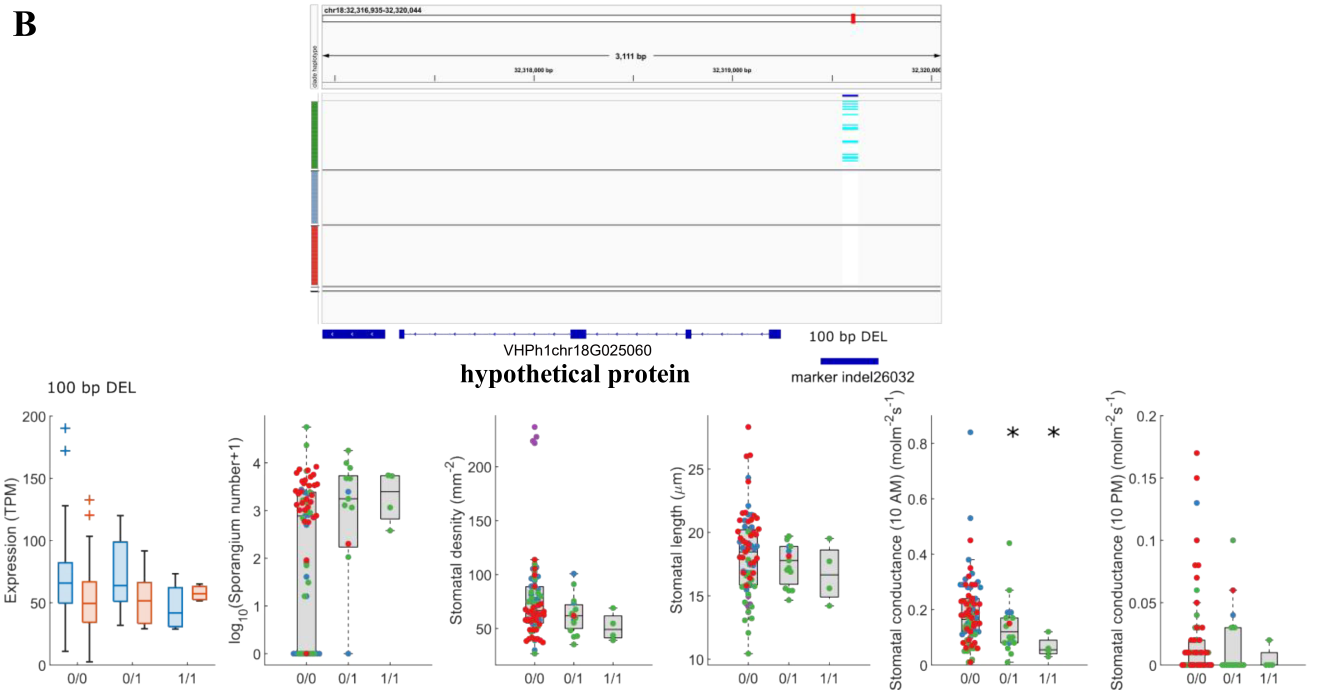

### Figure S10. Expression and phenotype of top genes in eQTL analysis

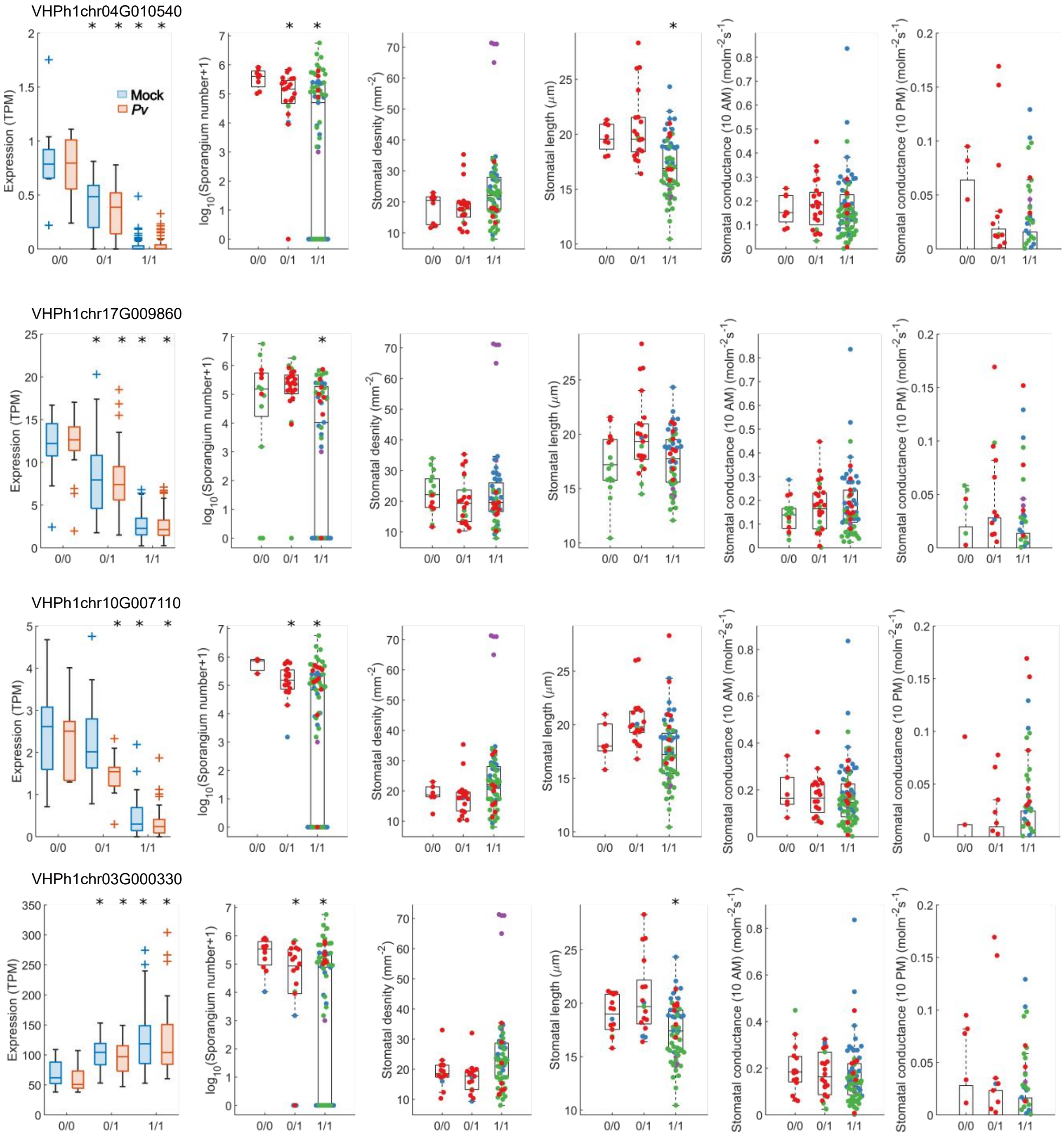
